# Discovering Fragile Clades And Causal Sequences In Phylogenomics By Evolutionary Sparse Learning

**DOI:** 10.1101/2024.04.26.591378

**Authors:** Sudip Sharma, Sudhir Kumar

## Abstract

Phylogenomic analyses of long sequences, consisting of many genes and genomic segments, infer organismal relationships with high statistical confidence. But, these relationships can be sensitive to excluding just a few sequences. Currently, there is no direct way to identify fragile relationships and the associated individual gene sequences in species. Here, we introduce novel metrics for gene-species sequence concordance and clade probability derived from evolutionary sparse learning models. We validated these metrics using fungi, plant, and animal phylogenomic datasets, highlighting the ability of the new metrics to pinpoint fragile clades and the sequences responsible. The new approach does not necessitate the investigation of alternative phylogenetic hypotheses, substitution models, or repeated data subset analyses. Our methodology offers a streamlined approach to evaluating major inferred clades and identifying sequences that may distort reconstructed phylogenies using large datasets.

## INTRODUCTION

Evolutionary biologists frequently assemble long sequence alignments containing numerous genes and genomic segments to resolve species relationships (Kumar et al. 2012; Kapli et al. 2020; Young and Gillung 2020; Kumar 2022). This advance has greatly increased the accuracy and resolution of inferred organismal relationships using phylogenomic methods (Rokas et al. 2003; Philippe et al. 2005; Edwards 2016; Williams et al. 2019; Homziak et al. 2023). However, despite using many-fold larger numbers of genes than needed to achieve high statistical significance theoretically (Rokas et al. 2003; Phillips et al. 2004; Gadagkar et al. 2005; Kumar et al. 2012), phylogenomic studies can produce species relationships that are not robust (Redmond and McLysaght 2021; Hughes et al. 2023). Dataset changes involving even a minute number of sequences have been reported to produce different evolutionary relationships (Phillips et al. 2004; Chiari et al. 2012; Salichos 2014; Smith et al. 2015; Brown and Thomson 2016; Shen et al. 2017; Shen et al. 2021). For instance, the exclusion of a single gene among 1,233 was associated with the unstable placement of a fungus family (Shen et al. 2017), and one exon was reported to destabilize highly supported clades inferred from an entire phylogenomic dataset (Smith et al. 2020). Such genes and sequences may bias the results because they are contaminants, such as paralogs, and/or the substitution models used do not adequately model gene- or species-specific molecular evolutionary dynamics (Chiari et al. 2012; Feuda et al. 2017).

Overall, such results challenge the intuition that the cumulative phylogenetic signals from many genes will neutralize the effects of a few outlier sequences and model assumptions (Gadagkar et al. 2005; Abadi et al. 2019; Kapli et al. 2020; Young and Gillung 2020; Kumar 2022; Guimarães Fabreti and Höhna 2023). Instead, these outlier sequences can dictate phylogenies inferred from big datasets, a phenomenon becoming increasingly common (Jeffroy et al. 2006; Hughes et al. 2023; Steenwyk et al. 2023). This pattern likely results from the bias introduced by outlier sequences that persist and determine phylogenetic relationships, while the statistical variance decreases quickly with increasing genes and sites (Philippe et al. 2005; Kumar et al. 2012; Kapli et al. 2020). Some differences in species relationships inferred from the concatenation, consensus, and coalescent approaches in phylogenomics are also attributable to the effects of outlier sequences (Mirarab et al. 2014; Smith et al. 2015; Shen et al. 2017; Homziak et al. 2023; Hughes et al. 2023; Shao et al. 2023).

Researchers are keen on pinpointing gene-species combinations that may unduly impact phylogenetic inference from phylogenomic data matrices containing thousands of gene-species combinations. Identifying such combinations is akin to searching for a needle in a haystack when investigators have already tried to remove non-orthologous sequences (Struck 2014; Steenwyk et al. 2023). Current solutions typically rely on evaluating alternative phylogenies, but these are not designed to isolate individual gene-species combinations and require time-consuming iterative reanalysis of data (Brown and Thomson 2016; Shen et al. 2017; Walker et al. 2018). For instance, the difference in gene-wise maximum likelihood (ML) support for alternative phylogenetic hypotheses has been used to rank influential genes and repeated phylogenomic analyses excluding the most discerning genes used to test their effect; see a review in (Steenwyk et al. 2023). This process necessitates a prior selection of clade to investigate as well as the knowledge of plausible alternative phylogenetic hypotheses and substitution models. However, only a limited set of clades or hypotheses may be testable in this type of analysis due to the lack of prior knowledge or an excess of plausible combinations. In addition, repeated ML and Bayes factor analyses impose a substantial computational burden (Liu et al. 2011; Höhna et al. 2021).

Instead of alternative phylogenies and substitution models, some approaches analyze different subsets of genes and species to look for fragile clades in the phylogeny inferred from the entire dataset. For example, subsamples containing varying numbers of genes were analyzed to assess the stability of the placement of certain species in the inferred phylogeny (Song et al. 2012). However, choosing the optimal subsample size and determining the number of subsamples to analyze can prove challenging (Edwards 2016), and such efforts may not even reveal the gene-species combinations that cause clade fragility. While such limitations are common among methods designed to identify outlier genes (Brown and Thomson 2016; Shen et al. 2017; Walker et al. 2018; Koch 2021), a few approaches aim to detect outlier sequences (gene-species combinations) by analyzing inferred gene trees and reporting outlier sequences, for example, associated with spuriously large pairwise distances in the inferred gene trees (Vienne et al. 2012; Comte et al. 2023). However, these outlier sequences are not detected for specific clades, and identifying fragile clades requires additional analyses.

Here, we present a new approach that uses evolutionary sparse learning (ESL) to identify fragile clades and the associated gene-species combinations without conducting additional phylogenetic inference with data subsets, substitution models, or phylogenetic alternatives. In brief, the ESL approach builds a (regularized) regression model in which genes and sites are explanatory variables, and a taxon’s presence or absence (1 or 0, respectively) in the clade of interest is the outcome. In ESL, one parameter penalizes the inclusion of genes (λ_G_), and another penalizes the inclusion of sites (λ_S_) in the clade-specific genetic model. For the given pair of penalty parameter values, ESL evaluates a large combination of genes and sites to determine one that correctly classifies the member taxa of an inferred clade using the fewest variables (Kumar and Sharma 2021).

In our investigation of ESL models built using a range of penalty values, many models for a clade could not classify member taxa in the clade with high confidence. This observation was surprising because the counts of genes and sites greatly exceed the number of taxa in any clade in phylogenomic alignments. This observation led to the formulation of two new metrics. One is the gene-species concordance (GSC), which identifies gene-species combinations harboring concordant (GSC > 0) or conflicting (GSC < 0) phylogenetic signals for the clade of interest. The second is the clade probability (CP; 0 ≤ CP ≤1) derived from all the GSC values and intended to pinpoint fragile clades in the inferred phylogeny. The estimation and use of GSC and CP do not need alternative phylogenies, substitution models, or data subsets. Their calculation does not require any pre-training or cross-validations in conventional machine learning approaches because the focus is on building a clade-specific genetic model rather than developing a classification system for use with the data not included in the alignment (Schrider and Kern 2018; Tao et al. 2019; Suvorov et al. 2020). We also implemented all these metric calculations in an analysis pipeline and packaged them in a distribution called *DrPhylo* (Fig. 1). This distribution can be downloaded as a standalone program for use on the command line (https://github.com/ssharma2712/DrPhylo) or accessed via a graphical user interface hosted in the MEGA software (an early beta version accessible anonymously to reviewers at https://www.kumarlab.net/M12withDrPhylo).

**Figure 1.**
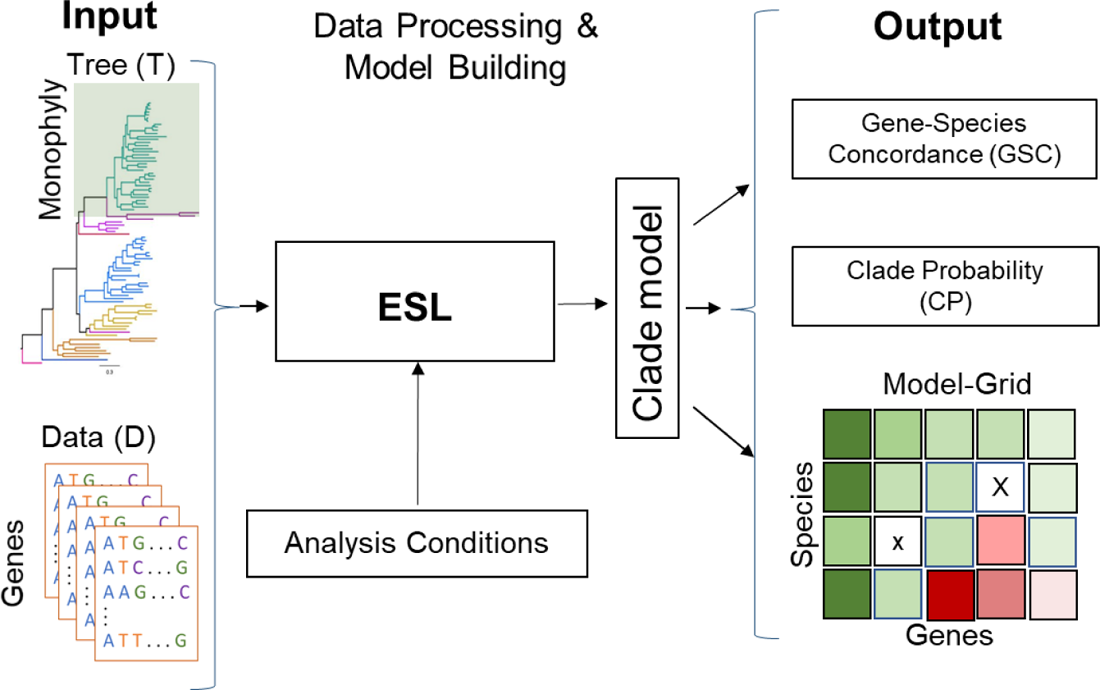
*DrPhylo* analysis pipeline. *DrPhylo* takes a phylogenetic hypothesis and a collection of FASTA files containing sequence alignments for individual groups of sites, e.g., genes, genetic segments, or any collection *of* sites (D). It is designed to accept the phylogenetic hypothesis in a text file (e.g., response.txt) or as a rooted phylogenetic tree with an identifier for the clade of interest in the tree written in the Newick format (T). These inputs are transformed into numeric data. Users specify options for *DrPhylo* analysis through the command line, including the range of the sparsity parameters. *DrPhylo* implements a phylogeny-aware class balancing, explained in the *Materials and Methods* section, builds the clade models for given sparsity parameter(s), and calculates metrics presented in this article. *DrPhylo* also outputs a graphical representation of the clade model in a grid format (Model-grid), which displays gene-species concordances (GSCs) and species classification probabilities (SCPs), as shown. *DrPhylo* also has a QUICK analysis option that employs a stopping rule to avoid building extremely sparse models containing genes less than a user-specified number (see *Materials and Methods*).

We used the standalone version of *DrPhylo* on a Linux computer to analyze multiple empirical phylogenomic datasets in which fragile clades and influential genes were previously reported (Wickett et al. 2014; Shen et al. 2016; Shen et al. 2017; Shen et al. 2018). This collection included a fungus dataset (86 species and 1,233 genes), an expanded fungus dataset (343 species and 1,292 genes), a plant dataset (103 species and 620 genes), and an animal dataset (37 species and 1,245 genes). Additionally, some clades in the inferred phylogeny are well-resolved with robust statistical support and unaffected by minor perturbations in the dataset. We used these datasets and species relationships as baselines to evaluate *DrPhylo*. Our analyses compared results from *DrPhylo* with other statistical approaches (e.g., ML and Bayesian) to gauge the effectiveness and efficiency of the new metrics in identifying overly influential and disruptive gene-species combinations and fragile clades.

## RESULTS

In the following, we first describe the approach for estimating GSC and CP using an example dataset of 1,233 nuclear gene alignments (609,899 amino acid positions) from 86 fungi species (Shen et al. 2016; Shen et al. 2017). The ML analysis of the concatenated supermatrix inferred clade A to be a sister to clade B (Fig. 2a). However, another phylogenomic study recovered an alternative phylogenetic placement for clade A, which was the sister to a group of clades B and C with very high (100%) bootstrap support (Riley et al. 2016). These two alternative hypotheses (Fig. 2b and 2c) for the placement of the clade A were previously compared by Shen et al. (2017) using ML analysis of 1,233 nuclear gene datasets. They reported a single gene to cause the fragility of A+B, which was the clade of interest (44 species) in the *DrPhylo* analysis.

**Figure 2.**
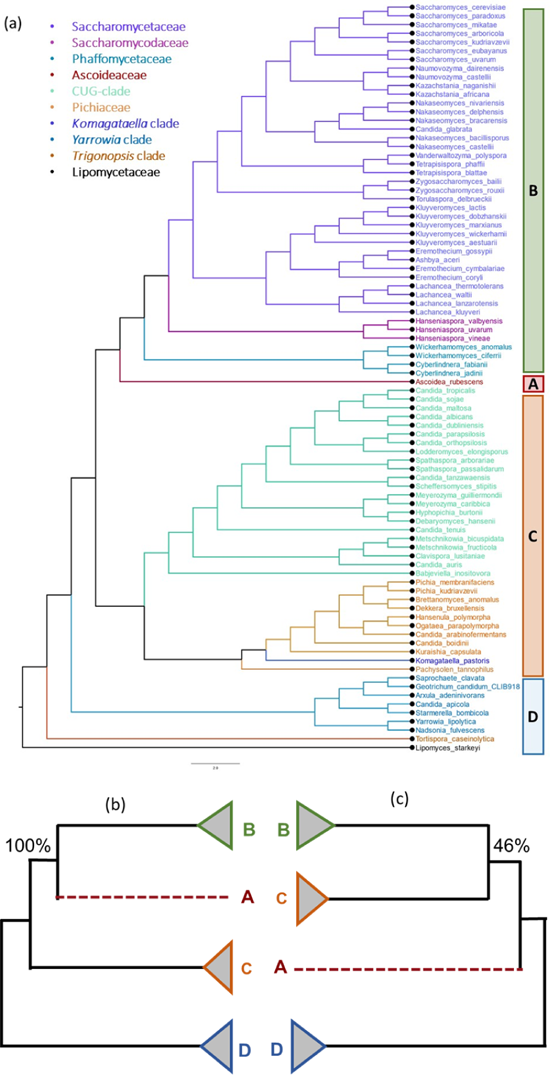
Contrasting phylogenetic relationships of three fungal clades. **a)** The maximum likelihood phylogeny of fungi inferred from a concatenated supermatrix of 1,233 nuclear genes (609,899 amino acid sites) by Shen et al. (2017). Clade A contains only *Ascoidea rubescence* (*Ascoideaceae*) and is sister to Clade B, which has 43 species of *Saccharomycetaceae*, *Saccharomycodaceae*, and *Phaffomycetaceae*. Clade C consists of 11 species of *Pichiaceae* and 22 CUG-Ser2 species (Shen et al. 2016; Shen et al. 2017). Clade D is the outgroup consisting of 9 species. Clade A+B received 100% bootstrap support in the concatenated supermatrix analysis (Shen et al. 2017). Contrasting evolutionary relationships of clades A, B, and C are shown in panels **b** and **c**, along with their bootstrap supports.

### Estimating Gene-Species Concordance (GSC)

In the first *DrPhylo* analysis, we built an ESL model for clade A+B, assuming a fixed pair of sparsity parameters for including sites and genes in the genetic model (λ_S_ = 0.1 and λ_G_ = 0.2, respectively). We will relax this assumption below. The A+B clade model included only 176 sites from 15 genes (see *Materials and Methods* for details of the options used). We expected sequences of these genes in all member species of clade A+B to harbor phylogenetic substitutions concordant with their placement inside A+B because the pattern-matching algorithm in sparse learning is expected to select optimal sites and genes at which the base configuration in the sequence alignment correlates with the presence of species in the clade A+B to the exclusion of the rest of the phylogeny.

We defined a gene-species concordance (*gsc*) metric to assess the degree to which a given gene in a given species harbors phylogenetic signals concordant with the clustering of taxa in A+B (see *Materials and Methods*). Biologically, we expected *gsc* values for all gene-species combinations to be positive for 15 genes included in the clade model. Instead, we found negative *gsc* values for many gene-species combinations, some of which were large in magnitude. The most extreme negative *gsc* value (−0.27) was for the gene *BUSCOfEOG7TN012* (*7TN012* hereafter) of *Ascoidea rubescens* (clade A).

To avoid reliance on an arbitrary choice of λ_S_ and λ_G_, we built 81 models for clade A+B using the range of site and gene sparsity parameters (0.1 ≤ λ_S_ and λ_G_ ≤ 0.9; step size = 0.1). Of these, only 12 models contained multiple genes and were retained for further analysis (see *Materials and Methods*). We defined *GSC* as the median *gsc* for a given gene-species combination across all multi-gene ESL models (see *Materials and Methods*).

Figure 3a shows the distribution of GSC scores for all gene-species combinations for clade A+B. In this distribution, two outlier GSC humps are seen. One on the right side (green, positive) involves the gene *BUSCOfEOG7W9S5*1 (*7W9S51* hereafter), which was the most influential gene identified previously (Shen et al. 2017). The hump on the left involves *7TN012* (red inset), which was not identified in any of the previous analyses (Fig. 3a). These two, and some other gene-species combinations, are easily visualized in a grid representation shown in Fig. 3b (M-grid for clade A+B). It quickly reveals that *7W9S51* provides the strongest phylogenetic signal (dark green) for placing all member species in clade A+B. In contrast, the gene *7TN012* carries the strongest conflicting signal (dark red) in the species *A. rubesence* (Fig. 3b).

**Figure 3.**
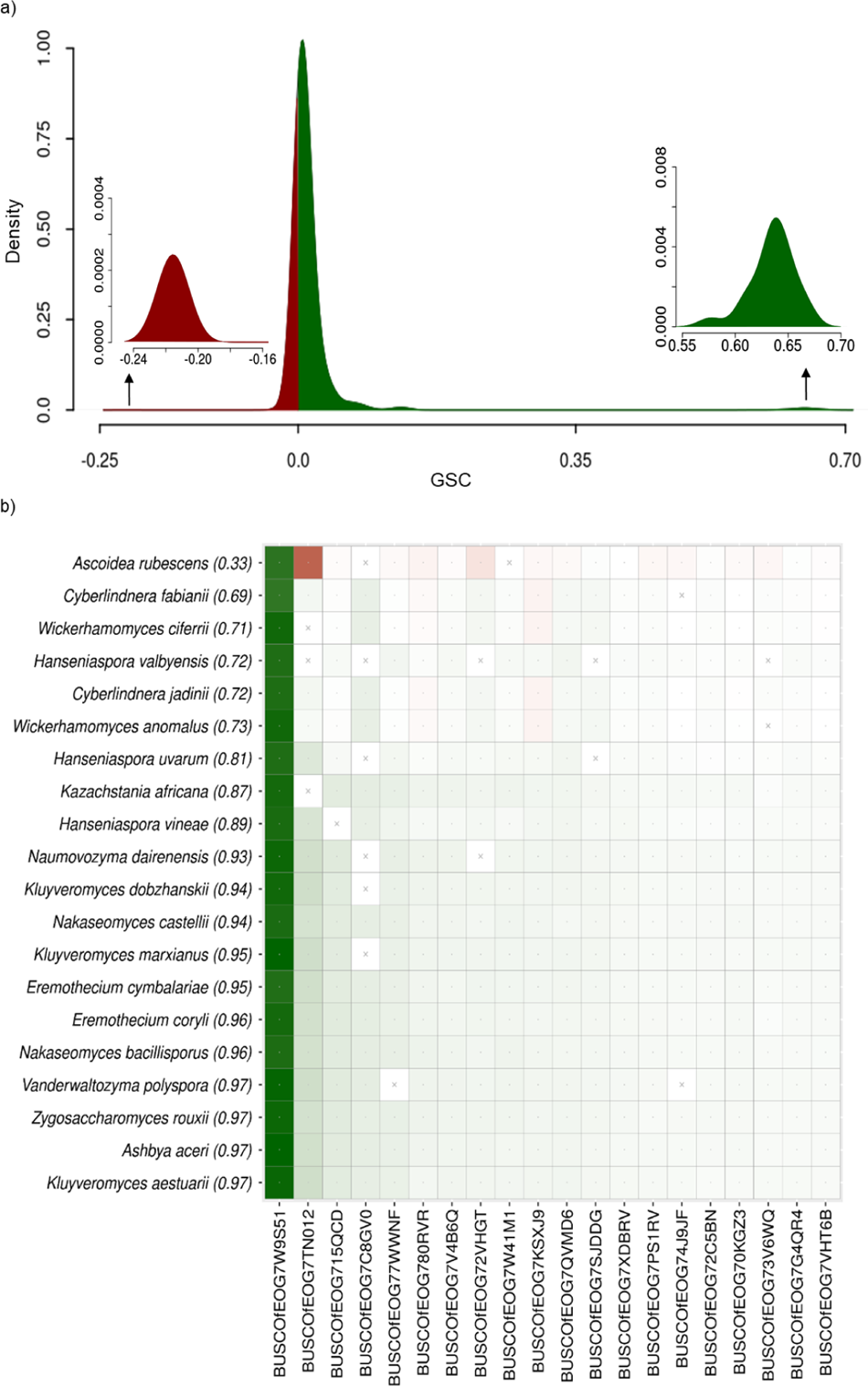
Distribution of GSC scores for clade A+B and the associated Model grid (M-grid). **a)** A histogram of *GSC* scores. The green inset on the right highlights gene-species combinations that show high concordance with the presence/absence of species in the evaluated clade. In contrast, the inset on the left (red) corresponds to negative *GSC* values and exposes combinations conflicting with A+B. **b**) A model grid for the A+B clade. The color intensity marks the degree of concordance (green) or discordance (red) of individual gene-species combinations. A cross-mark indicates missing data. The top 20 species (out of 44) and 20 genes (out of 126) are shown. Among these 20 species, one is from clade A at the top of the grid, and the other 19 are from clade B. On the top-left are species with the lowest *SCP* (shown in parentheses) and genes receiving the highest average |*GSC*| across all species.

### Estimating the Clade Probability (CP)

To compute CP, we first estimate the species classification probability (*scp*), a logit transformation of the sum of all *gsc* scores for the given species *s* for a given pair of λ_S_ and λ_G_ values (see *Materials and Methods*). To avoid reliance on a specific pair of parameter values, we computed *scp* from all 12 multi-gene models and then estimated a single SCP value for each member species of the clade of interest (*Materials and Methods*). Then, SCPs for all member species in a clade were used to estimate CP, which measures the robustness of the clade of interest. CP is simply the minimum of all SCPs. The CP of A+B is low (0.33) because the SCP of *A. rubescence* to be clustered with the clade B is low (SCP = 0.33).

### Further analysis of fungus relationships

We now present results from the full *DrPhylo* analysis of clade A+B in the above dataset, whose low CP (0.33) is in stark contrast with its high bootstrap support (100%) in the ML analysis of the concatenated supermatrix (Shen et al. 2016; Shen et al. 2017). The low CP is caused by *7TN012*, and some other genes, that do not support this grouping (GSC < 0; Fig. 3b). The negative GSC score for *7TN012* is well-justified by its gene tree, in which *A. rubescence* (clade A) is positioned far from clade B, indicating gene tree-species tree discordance (Supplementary Fig. 2). However, Bayes Factor (BF) analysis using alternative hypotheses (Fig. 2b-c) did not find *7TN012* to be unusual, as it ranked 938 out of 1,233 genes based on 2*ln*(BF). Also, the role of *7TN012*, which is a homolog of the GLT1 gene in *Saccharomyces cerevisiae*, was not revealed in the ML analysis of these alternative hypotheses (Shen *et al*. 2017). Also, PhylteR, an outlier detection approach using multi-dimensional scaling (Comte et al. 2023), did not identify any gene-species combinations involving *7TN012* in its output of 681 outlier sequences. It is likely because PhylteR analysis is not focussed on the clade of interest. However, PhylteR finds *7W9S51* to be an outlier, but it does not indicate whether it is supportive or disruptive of the inferred phylogeny.

We also used the approximate unbiased test (AU-test) to compare the species tree (Fig. 2a) with the gene trees for *7W9S51* and *7TN012.* We expected that the *7W9S51* gene tree would be concordant with the inferred global phylogeny but not *7TN012*’s gene tree. Surprisingly, the AU-test rejected the inferred global phylogeny for both gene alignments (*P* < 0.05). Similar results were obtained for other influential genes identified in the *DrPhylo* analysis (Supplementary Table 1).

These findings indicate that *DrPhylo* can complement conventional statistical methods by offering insights into highly influential and conflicting gene-species combinations associated with the fragile clade.

### Impact of influential genes and gene-species combinations on inferred phylogenies

The M-grid reveals that the placement of *A. rubescence* in clade A+B is fragile, receiving the strongest support from *7W9S51* (GSC = 0.58), while a majority of the genes (68%) in *A. rubescence* contradict the grouping of A and B clades (GSC < 0 in the M-grid; Fig. 3b).

Therefore, the removal of *7W9S51*, with large positive GSC, may decrease the support for A+B, while the removal of genes with negative GSC may do the opposite. However, the impact of such removals on the final phylogeny produced by the concatenation matrix analysis is not easily predictable in our experience because the biases caused by the remaining genes cannot be anticipated *a priori*.

In any case, the hypothesis that excluding *7W9S51* would reduce the support for the clade A+B was tested previously, and the reduced dataset united clade B with clade C rather than A (Shen et al. 2016; Shen et al. 2017). The bootstrap support for A+B was reduced to 54% from 100% for the whole dataset (Fig. 2c). The bootstrap support did not decline (61%) after the subsequent removal of the *7TN012* gene. This fragility was also evident from the MSC analysis before and after excluding *7W9S51*, *7TN012*, or both, as the posterior probability for clade A+B was low in all cases (64% - 68%) because the MSC approach is resilient to the exclusion/inclusion of one or a few genes in the dataset (Shen et al. 2017; Mirarab et al. 2021; Warnow 2015).

Overall, the low bootstrap support and conflicting placement for clade A after the removal of a few genes established the fragility of the clade A+B, which *DrPhylo* could successfully identify along with associated genes without needing to perform phylogenetic analyses with data subsets or alternative evolutionary hypotheses. Once these genes are identified, one can inspect their gene trees, which we did for *7W9S51* and *7TN012*. We found an unusually large separation between clade A+B and other species (5.86 substitutions per site) in the *7W9S51* gene tree (Supplementary Fig. 1). This long branch likely amplifies the phylogenetic information favoring clade A+B in the concatenation analysis. Consequently, excluding *7W9S51* from the dataset significantly reduces support for A+B. In contrast, clade A+B is not monophyletic in the *7TN012* gene tree (Supplementary Fig. 2).

### ESL analysis of an expanded fungus data set

Shen *et al*. (2018) collected data from three additional species for clade A, one member of *Ascoideacea* and two species of *Sacchromycopsis*, to re-examine the evolutionary relationships among *Fungi*. The number of species was also increased in clade B as well as other clades, and the number of genes was increased to 1,289. However, despite data expansion, the M-grid for A+B (Fig. 4) did not show an increase in *CP* for this clade A+B. Rather, it decreased to 0.00 due to low *SCP* for two newly added *Sacchromycopsi* species. *DrPhylo* found clade A+B fragile because 49% of the *GSC* values were negative for these *Sacchromycopsi* species from clade A. This result is consistent with low quartet support of 39% and gene concordance factor (gCF = 19.6%) for A+B. Interestingly, the clade A+B is recovered with high statistical support (100%) in both concatenation and MSC approaches with or without *EOG09343FGH*, making it an enigmatic dataset for resolving the relationship of A, B, and C.

**Figure 4.**
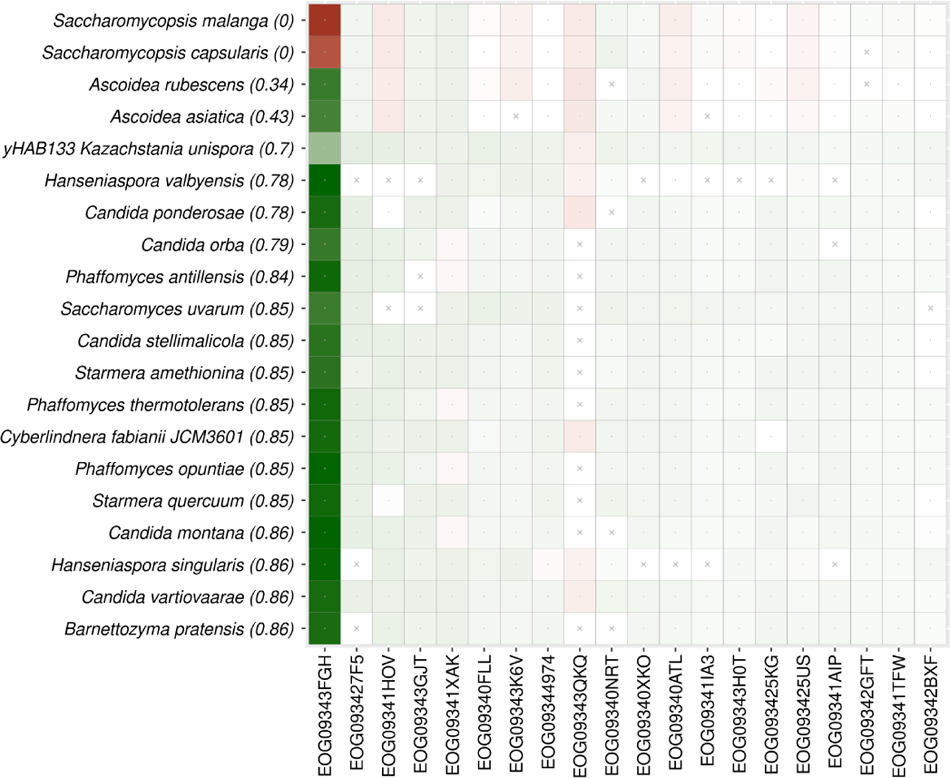
An M-grid from the extended fungal dataset. The M-grid for clade A+B for the extended fungus dataset. The top four rows in the grid comprise species belonging to clade A, while the remaining species are from clade B The color intensity marks the degree of concordance (green) or discordance (red) of individual gene-species combinations. A cross-mark indicates missing data. The top 20 species and 20 genes are shown. On the top-left are species with the lowest *SCP* (shown in parentheses) and genes receiving the highest average |*GSC*| across all species.

*DrPhylo* identified *EOG09343FGH* to harbor the strongest contradictory phylogenetic signals (Fig. 4). Notably, *7W9S51* in the previous dataset and *EOG09343FGH* are homologs of the DMP1 gene in the model system *Saccharomyces cerevisiae* (Shen et al. 2017; Shen et al. 2018). An inspection of the *EOG09343FGH* gene tree (Supplementary Fig. 3) revealed the same problem as *7W9S51*, i.e., its gene tree contains an unusually long internal branch (6.2 substitutions per site). In addition, two *Saccharomycopsis* species of clade A are on the opposite ends of this branch. That is, clade A was not monophyletic, and some of its member species have far greater sequence divergence from each other than with members of other clades. Such gene tree patterns may arise due to hidden paralogy or other biological factors, such as horizontal gene transfer, a frequently observed phenomenon in many clades of fungal species (Richards et al. 2009; Schmitt and Lumbsch 2009; Fitzpatrick 2012; Shen et al. 2018; Steenwyk et al. 2023). Further the ML analysis of two alternative hypotheses for A+B also detected *EOG09343FGH* as having the highest likelihood difference, and PhylteR identified *EOG09343FGH* as containing the largest number of outlier sequences (338 out of 1,260). However, PhylteR’s outliers are not tied to specific clades.

In summary, *DrPhylo* successfully pinpointed conflicting gene-species combinations involving *Sacchromycopsis* species and the *EOG09343FGH* gene without needing gene phylogenies, substitution models, or alternative species relationships for clade A+B.

### DrPhylo analysis of a “control” fungus clade

In addition to analyzing the known fragile clades above, we tested new metrics on a 36-species clade of *Saccharomycetaceae* that was used as a control in a previous study to validate the ML analysis approach (Shen et al. 2017). For this clade, *DrPhylo* analysis produced a model in which all the GSCs were positive, i.e., they harbored phylogenetic signals concordant with the monophyly of the clade analyzed. The M-grid for this comparison is shown in the Supplementary Figure 4. The CP for this clade was high (0.79), confirming the results from the ML analysis.

We also used the data analyzed in the above analysis to investigate the ability of *DrPhylo* to detect outlier gene-species combinations in synthetic datasets in which we deliberately introduced introgression across species in the most important gene *BUSCOfEOG715QCD* (see Supplementary Figure 4-5). *BUSCOfEOG715QCD* is an ortholog of the SPT6 gene (*YGR116W*) in *Saccharomyces cerevisiae*. We generated 100 such datasets by swapping the selected gene sequences between two randomly selected species, one from the *Saccharomycetaceae* clade and the other from outside the clade. Because the errors were introduced in the most important gene, we expected this gene to be included in the ESL model and the affected gene-species combinations to receive negative GSC values.

In *DrPhylo* analyses, *GSC* was indeed negative for the affected gene-species combinations in 98 synthetic datasets and was positive but close to zero for the other two (Fig. 5a). That is, *DrPhylo* showed 98% accuracy in detecting errors in the most influential genes. A similar performance (98%) was observed when the introgression was one-way in which a randomly selected *Saccharomycetaceae* species received the gene sequence from a randomly selected species outside the *Saccharomycetaceae* clade (Fig. 5b). In this case, CP was relatively high for all the *Saccharomycetaceae* clades in all the synthetic datasets (0.88-0.93), showing that the phylogenetic inference is robust despite the presence of some data errors. This pattern is likely because the stem branch for this clade is ∼10 times longer than that for clade A+B in the fungi phylogeny, presented in the first example.

**Figure 5.**
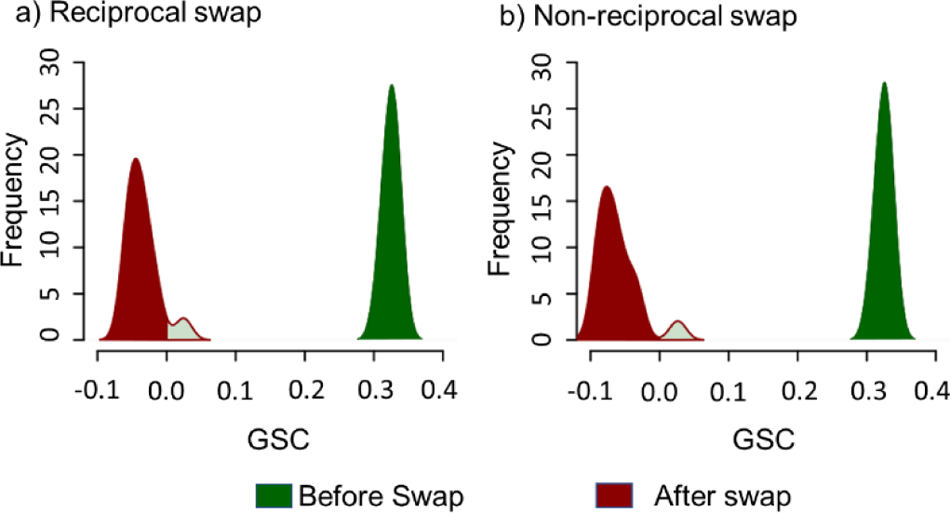
Gene-Species Concordance (GSC) Scores of Simulated Errors. The change in *GSC* scores for gene-species combinations with (a) reciprocal and (b) non-reciprocal swaps. Before the swap, their GSC scores were positive (green density plots). After the swap, GSC scores became negative (red density plots). Mild green with a red border in the red density indicates cases in which the simulated errors were not detected.

We also applied PhylteR to these simulated datasets, which produced many outliers for every dataset, including *7W9S51* and the gene *BUSCOfEOG715QCD* that underwent introgression between species. Neither ML nor *DrPhylo* analyses found *7W9S51* to be influential for this control clade, but the PhylteR diagnosis is not clade-specific, so the outliers reported are for the whole phylogeny.

### Analysis of a phylogeny of plants

To assess the generality of the results presented above for *DrPhylo* analysis of the fungus dataset, we applied *DrPhylo* to the phylogeny inferred in an analysis of 620 nuclear gene sequences from 103 plant species in which the focus was on identifying the closest relatives of *Chloranthales* (C). The concatenated supermatrix approach united *Eudicotidae* (E) and *Chloranthales* with a bootstrap support of 100% for C+E (Supplementary Fig. 6) (Wickett et al. 2014; Shen et al. 2017). *DrPhylo* found C+E to be fragile, as the CP was low because of *Saracandra glabra* (SCP = 0.37). *S. glabra,* the only member of clade C, received low SCP because 33.9% of genes (97 out of 286) did not support its placement inside clade C+E. The M-grid for this clade revealed some influential genes (e.g., *6040_C12*, *4490_C12*, *4478_C12*) that strongly support the clustering of C with E.

The gene *6040_C12* (orthologues of *AT3G46220* gene in *Arabidopsis thaliana*) has the highest influence in placing *S. glabra* (C) in the clade C+E (Fig. 6). The *6040_C12* sequences in five species in clade E harbor conflicting phylogenetic signals (red cells, Fig. 6) for the clade C+E. These five species grouped far away, separated by a long internal branch, 0.8 substitutions per site, from other members of the C+E clade in the *6040_12* gene phylogeny (Supplementary Fig. 7). Two other genes, *4478_C12* (ortholog of *AT4G02580* gene in *A. thaliana*) and *4490_C12* (ortholog of *RbcX2* gene in *A. thaliana*), received negative GSCs in the same five species similar to *6040_12*. Their gene trees showed patterns similar to the *6040_C12* gene tree, including a long branch length separating the same five species of C+E from the rest. There was a large effect of *6040_C12* on the phylogeny produced from the concatenated supermatrix of 619 genes that excluded *6040_C12*. The ML phylogeny united *Chloranthales* with *Magnolids* (C+M) with 71% bootstrap support, which is different from the whole dataset analysis that produced C+E with high support. The species tree inferred from the MSC approach before and after the removal of *6040_C12* assigned a low posterior probability of 0.30 to C+E in both analyses, as C+M received a 57% local posterior probability (Shen et al. 2017). By the way, these patterns are consistent with previous reports that used two alternative phylogenetic hypotheses about the placement of Chloranthales in the ML analysis (Shen et al. 2021).

**Figure 6.**
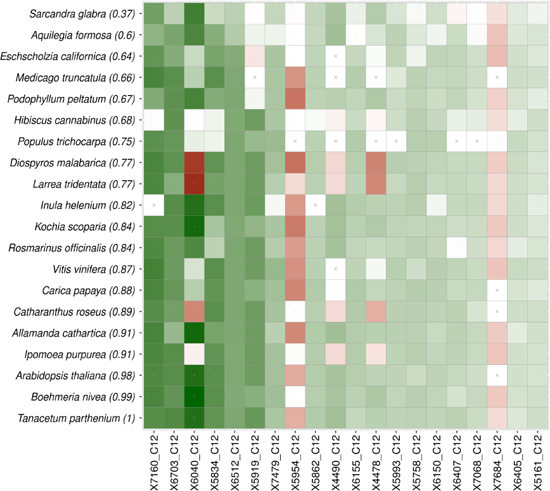
The M-grid for clade C+E in the phylogeny of plants. The M-grid for the C+E clade contains 20 species from the plant phylogeny. A total of 20 genes (out of 292) are displayed and ordered using the average positive *GSC*. The color intensity marks the degree of concordance (green) or discordance (red) of individual gene-species combinations. A cross-mark with a white background indicates missing data. The top 20 species and 20 genes are shown. On the top-left species with the lowest *SCP* (shown in parentheses) and genes receiving the highest average |*GSC*| across all species. The species on the top-left is from clade C, and the other 19 species are from clade E.

In addition to *6040_C12*, the M-grid reports two additional genes, *5954_C12* and *7684_C12,* to be not supportive of clade C+E (Fig. 6). Their gene trees do not have a C+E clade, as C and E are located distantly in the phylogeny (Supplementary Fig. 8 and 9). The PhylteR analysis of this dataset also found *6040_C12* but not *5954_C12* and *7684_C12. PhylteR* reported additional genes (*4478_C12* and *4490_C12*) that may impact other clades in the inferred phylogeny. Therefore, new metrics successfully identified the fragile clade (C+E), problematic species (*S. glabra*), and influential as well as disruptive outlier sequences.

### Analysis of an animal phylogeny

Finally, we applied *DrPhylo* to a phylogeny of 37 rodents inferred from a phylogenomic dataset of 1,245 nuclear genes. The ML phylogeny inferred from the concatenated supermatrix places *Pogonomelomys ruemmleri* (P) outside of the SHL clade (see Supplementary Fig. 10) with a high rapid bootstrap support (98%). *DrPhylo* produced a low CP (0.66) for the SHL clade (Fig. 7), designating it a fragile clade with three of the member species receiving low SCP scores (0.66 - 0.67). About 40% of the genes in these three species received negative GSCs in the clade model (Fig. 7). None of these genes were identified in the ML analysis of alternative hypotheses or by PhylteR, even though SHL clade is not monophyletic in their phylogenies (Supplementary Fig. 11). However, the fragility of SHL was observed in MSC analysis, which inserted *P. ruemmleri* inside the SHL clade with a high posterior probability (LPP = 95%).

**Figure 7.**
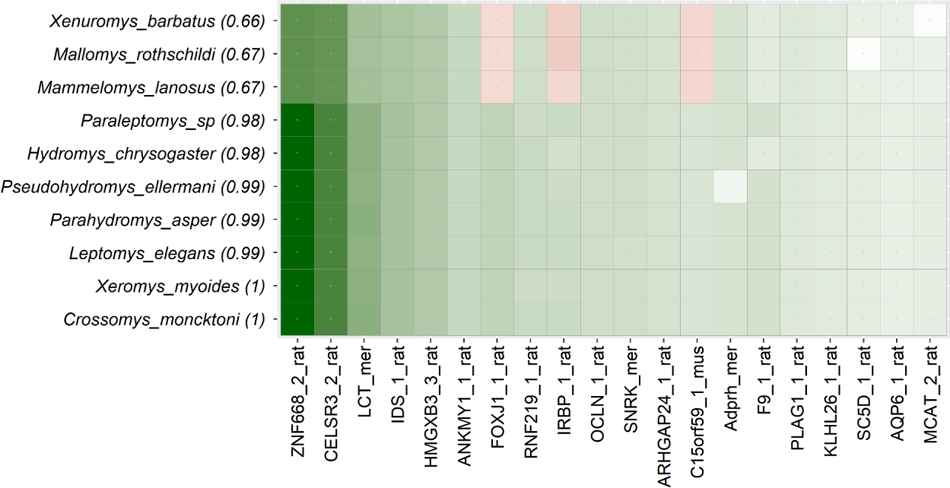
The M-grid for clade SHL. The M-grid for the SHL clade shows 20 genes (out of 260). These genes are ordered using the average of absolute GSC. The color intensity marks the degree of concordance (green) or discordance (red). All of these species (10) were selected from the SHL clade using smart sampling to balance the clade of interest and the outside world.

The fragility of the monophyly of the SHL clade, as well as the placement of *P. ruemmleri*, was not attributed to a few genes or sequences (Shen et al. 2021) but likely resulted from incomplete lineage sorting (Roycroft et al. 2020). The ML analyses identified a few other genes (*LCT_mer* and *IDS_1*) to be highly influential, which exhibit strong support for the SHL clade as shown in the M-grid (Fig. 7). Previously, the exclusion of these genes did not alter the inferred phylogeny and the SHL clade in the ML analysis of the concatenated sequence alignment (Shen et al., 2020). A previous study found 36% (451 out of 1,245 genes) of the total genes inconsistent between a pair of species tree hypotheses (Shen et al. 2021). The inferred species tree using the MSC approach after the removal of these genes became concordant to the ML tree from the concatenated sequence alignment (Supplementary Fig. 10).

In summary, *DrPhylo* could identify fragile clades that exhibit incongruence between the concatenation and MSC approach based on the analysis of the inferred phylogeny alone.

## Conclusions

We have advanced the use of evolutionary sparse learning (ESL) to diagnose phylogenetic instability and likely causal genes and species through novel metrics that detect fragile clades and underlying gene-species combinations. We have established the utility and abilities of ESL models and these metrics using empirical and synthetic datasets. The use of new metrics is made practical by the computationally efficient tool *DrPhylo,* which required less than 30 minutes for the analysis of the smaller fungus dataset (86 species, 1233 genes, and 609,899 sites) and 52 minutes for the expanded dataset (343 species, 1292 genes, and 527,069 sites); (see Supplementary Table 2). This means that *DrPhylo* can quickly scan major clades of the inferred phylogenomic tree without requiring the knowledge of problematic clades or alternative phylogenetic hypotheses. *DrPhylo* will reveal individual sequences (gene-species combinations), which we have shown to produce novel findings in analyzing three empirical datasets. In *DrPhylo*, an investigator may partition the data based on any desired biological annotations, including genes, proteins, codon positions, exons, and functional elements. Also, groups of sites can be inferred using statistical approaches that partition the data into evolutionarily homogeneous segments (Yang 1996; Kumar et al. 2012; Lanfear et al. 2017). Every site in the alignment can belong to its group, which would be useful when the data consists of only one gene or genomic segment.

*DrPhylo* does not necessitate in-clade phylogeny or conduct ML calculations using a base substitution model. Therefore, the identification of fragile clades and causal sequences (gene-species pairs) are agnostic to selecting a substitution model or any phylogenetic tree error within the clade of interest. *DrPhylo* also estimates signed concordance scores for each sequence, revealing which genes support or oppose species placement within the clade. While PhylteR and similar approaches also detect outlier sequences, these outliers are not clade-specific, as mentioned earlier. So, they require further analyses to determine which clades might be impacted by these outliers. Furthermore, the use of inferred gene trees makes the identification of outlier sequences susceptible to gene tree estimation error, a common challenge for methods using estimated gene trees.

We anticipate the new metrics presented here to be especially beneficial when only a small fraction of gene-species combinations carry signals that conflict with the placement of member taxa inside the clade of interest. This is because the ESL process of building clade models is unlikely to select genes whose sequences harbor phylogenetic signals conflicting with the membership of many species inside and outside the clade of interest. Therefore, if a gene with a significant amount of phylogenetic information for uniting species in the given clade has a limited number of disruptive gene-species combinations, then that gene will likely be included in the ESL models. Such sequences will receive negative GSC values in some genetic models and be recognizable as outliers in the M-grid. It is also advisable to apply *DrPhylo* for clades with a substantial number (e.g., ≥ 5) of taxa in the clade of interest, as machine learning methods generally demonstrate better performance for datasets with a large number of samples (e.g., taxa). Therefore, we suggest applying the new approach to well-curated phylogenomic datasets, like those analyzed here, to diagnose fragile clades and associated gene-species combinations following phylogenetic inference. While the gene-species combinations revealed in *DrPhylo* analyses may not always result in the fragility of the inferred clades, they are inherently intriguing, potentially stemming from biological processes such as gene losses and gains, introgression, and horizontal gene transfers (Chiari et al. 2012; Nakhleh 2013; Brown and Thomson 2016; Steenwyk et al. 2023).

## Acknowledgments

We thank Drs. Alessandra Lamarca, Jack Craig, Jose Barba-Montoya, and Xinghua Shi for reading the manuscript and providing many helpful suggestions. We thank Maxwell Sanderford for extensive technical support. We also thank Dr. Xing-Xing Shen and two anonymous reviewers for their thoughtful comments. This work was supported by a research grant from the National Institutes of Health to SK (R35GM139540-04).

## Data and codes availability

All sequence alignments and phylogenetic trees used in this article were obtained from published articles and have been assembled in a Figshare repository: https://figshare.com/s/590f73e9422d7dca0076. A GitHub repository containing scripts and software used to perform *DrPhylo* analyses and build model grids is available at https://github.com/ssharma2712/DrPhylo.

## Materials and Methods

### Evolutionary Sparse Learning (ESL)

An ESL model is defined as *f*(Y) = X*β*, where *f*(Y) is a logit link function of the category assigned to each species: +1 for member species of the clade of interest and −1 for all others in the given phylogeny (Kumar and Sharma 2021). In the ESL model, X is a one-hot encoded sequence alignment matrix produced as previously described (see Figure 1 in ref. (Kumar and Sharma 2021)). *β* is a column matrix of coefficients, estimated using bi-level sparse group LASSO regression that minimizes the logistic loss by penalizing the inclusion of individual sites (site sparsity parameter, λ_S_) and groups of sites such as genes (group sparsity parameter, λ_G_) to avoid model overfitting (Tibshirani 1996; Meier et al. 2008; Kumar and Sharma 2021). Groups can be collections of contiguous sites (e.g., genes, exons, introns, and proteins) or non-contiguous sites (e.g., codon positions), and sites with functional annotations (e.g., coding genes and non-coding elements), among other possibilities. Grouping sites based on biological and sequence features makes the ESL modeling a partitioned analysis common in phylogenomic studies (Hillis and Bull 1993; Mirarab et al. 2014; Kainer and Lanfear 2015).

In ESL, quantitative models with *β* estimates capture the strength of association between the pattern of sequence evolution at individual sites and genes with the presence and absence of species in the clade of interest. Generally, many genes and sites received a *β* value of 0 in the selected genetic model, leading to a sparse solution for clade-specific genetic models. ESL with bi-level sparsity differs from the contemporary machine learning approaches in ecology and evolution, which focus on classification by training machine learning models using synthetic data.

We transformed species relationships into a binary response (Y) and assigned +1 for all species in the monophyletic clade and −1 for species outside of the clade. Such binary classification is common in supervised machine learning of binary classification using the perceptron algorithm (Freund and Schapire 1999). Each gene sequence alignment was numerically transformed into binary one-hot encoded matrices (Kumar and Sharma 2021) and used as independent variables (X) for model building.

The myESL software, an open source library written in C++, implements ESL (Sanderford et al. 2024), which we used as the base for developing the pipeline for *DrPhylo* (https://github.com/ssharma2712/DrPhylo) for practical application of the methods and metrics presented here (Fig. 1).

### Building a Clade Model

*DrPhylo* first built many genetic models using the ESL approach that employed generalized least absolute shrinkage and selection operator (LASSO) logistic regression (Kumar and Sharma 2021). As the data are partitioned into groups of sites (e.g., genes) and we aim to select the highly influential genes and sites from genes, we used bi-level sparse group logistic LASSO regression. The ESL implementation applies the Moreau–Yosida Regularization algorithm (Liu and Ye 2010; Liu et al. 2011; Kumar and Sharma 2021) with 100 iterations (default) for convex optimization of the regression coefficients (β) for building the clade model.

### Estimation of Gene-Species Concordances (GSCs) and Clade Probability (CP)

For each clade model, we calculate the gene-species concordance (*gsc*) metric using the given ESL model to assess the degree of the concordance for a given gene (*g*) in a species (*s*), which is given as follows:

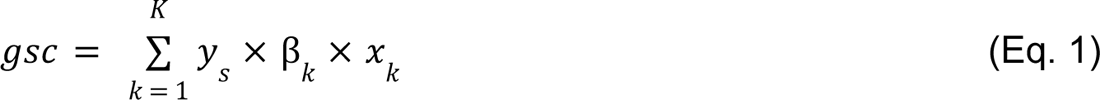

Here, *gsc* is the sum of the product of one-hot encoded bases (*x_k_*) of site *k* in the given gene *g* from species *s* with the numeric response for the species *s* and the regression coefficients (βs) in the ESL model. *K* is the number of one-hot encoded bases in the gene *g*. *gsc* quantifies the strength and direction of concordance. It is analogous to the SHAP value (Lundberg et al. 2020) to quantify a feature’s contribution to the predictive ability of a machine learning model. However, unlike SHAP, *gsc* does not require re-running ESL by excluding/including genes or sites in the model-building process.

We also calculate the species classification probability (*scp*) for each member species from each clade mode. The *scp* is the sum of all *gsc* and the model intercept (β0; equation 2 below)

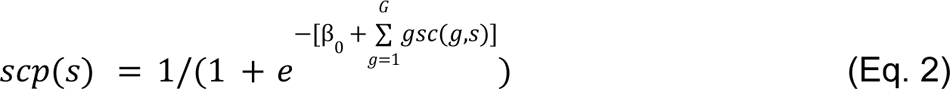

Here, *G* is the total number of genes in the dataset. This metric is the same as the standard classification probability in LASSO (Liu et al. 2011; Hastie et al. 2015). We normalized the *scp* for all member taxa to transform this metric to range from 0 to 1 for the given clade as follows:

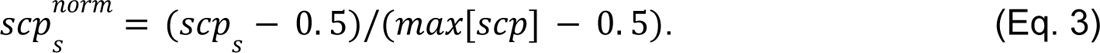

In this context, *scp_s_* denotes the probability of classification for a species *s*, while *scp* represents the array of probabilities encompassing all member species. We adopted a minimum *scp* of 0.5 since the predicted response for any member species, as determined by the clade model, is anticipated to be no less than zero. Therefore, a species with the minimum predicted response would receive a *scp* equal to 0.5. If the predicted response for a species is less than zero, then the clade model has misspecified the species. We set the normalized *scp* for those species to be zero.

### Gene Species Concordance (GSC) and Clade Probability (CP)

The GSC is the final estimate for the gene species concordance estimated by summarizing *gsc* from each genetic model built by a pair of the sparsity parameters. We ensembled all *gsc* values using a summary statistic, median. Mathematically, we define GSC for the given gene *g* and species *s* as follows.

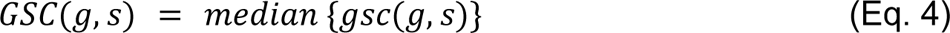

Here, {*gsc*(*g*, *s*)} is the vector of all *gsc* scores for gene *g* and species s estimated from ESL models. After normalization, we also summarized scp(*s*) for the species *s* from all ESL models to estimate the classification probability of ensembled species and defined them as *SCP(s)*. *SCP(s)* is the mean of all *scp(s)* for species *s* and is mathematically defined as follows.

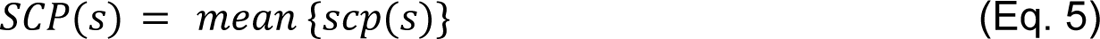

Here {*scp*(*s*)} is the vector of all *scp* scores for the species *s*. The clade probability (CP) for the clade of interest is the minimum of SCP from all member species and is defined as follows.

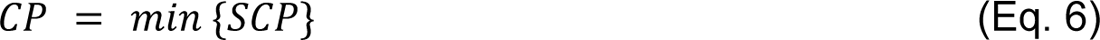

Here {*SCP*} is a vector of species classification probability estimated from the ensemble ESL model.

### Phylogeny-Aware Class-Balancing for ESL

To build an ESL model, we select species by phylogeny-aware class-balancing in which an equal number of species inside and outside the clade of interest were selected. When many outgroup species are available, then the closely related species are selected. For example, a given rooted phylogenetic tree with S_All_ species contains S_+1_ and S_-1_ species inside and outside the clade of interest, respectively; S_All_ = S_+1_ + S_-1_. To balance the number of species inside and outside the clade, we employed phylogeny-aware sampling when S_+1_ < S_All_/2 (S_+1_ < S_-1_; scenario 1) or S_+1_ > S_All_/2 (S_+1_ > S_-1_; scenario 2). In scenario 1, we first select clades from the outside +1 group that is the closest sister of the monophyletic clade of interest (+1 group) until S_+1_ ≤ S_-1_. If S_+1_ < S_-1_, we compute the pairwise distance between species (leaf nodes) in the S_-1_ set and remove one sequence randomly from the pair with the lowest distance. Next, one randomly selected species is removed from the pair with the second lowest pairwise distance, and this process is iterated until S_+1_ = S_-1_. Similarly, we sample species from the clade of interest in scenario 2 (S_+1_ > S_-1_) until S_+1_ = S_-1_.

### DrPhylo’s Quick option

We found that the number of genes included in the ESL model generally decreased monotonically with the site (λ_S_) and gene (λ_G_) sparsity parameters (Supplementary Fig. 12a-b), so we developed a simple stopping rule to avoid calculating models that will contain only one gene. *DrPhylo* begins with λ_S_ = 0.1, builds an ESL model starting with λ_G_ = 0.1, and counts the number of genes selected in the model. Then, λ_G_ is increased by the user-provided step size (Δλ; 0.1 by default) to build the next model, where λ_S_ is fixed. This process is stopped when the ESL model contains only one gene or λ_G_ becomes 0.9. This procedure provides an upper limit on λ_G_, i.e., λ_G,max_. In the next step, λ_S_ is increased by Δλ, and then models are built until λ_G_ reaches λ_G,max_. This process is repeated by increasing λ_S_ until a model contains only one gene. Then, all the models containing one gene are discarded before estimating the GSC and CP metrics described below.

### Data sets analyzed

#### Empirical datasets

Four empirical datasets were obtained from previous studies, which represent three major groups in the Tree of Life: Fungi, Plants, and Animals. Some species relationships in the inferred phylogenies from these datasets are known to be fragile because of highly influential outlier genes. The first fungus dataset, consisting of 1233 nuclear genes derived from 86 yeast species, was previously described by (Shen et al. 2017)). The length of genes in this dataset varied between 167 and 4854, and the number of taxa in each gene ranged from 39 to 86. The other taxon-rich fungus dataset comprised 343 yeast species and 1,292 nuclear genes and was analyzed by (Shen et al. 2018) *et al*. (2018). The plant dataset encompassed DNA sequences of 620 nuclear genes from 103 plant species (Wickett et al. 2014; Shen et al. 2017). The gene sequence alignments in this dataset were 6 to 1820 base pairs long and containing 55 to 103 plant species. The animal dataset contained 1,245 nuclear gene sequences from 37 rodent species. The number of species in each gene sequence alignment varied between 32 to 37, and the gene alignment lengths ranged from 249 to 7,413.

#### Synthetic Datasets with Simulated Contaminations

We introduced data errors in empirical datasets to assess the performance of new metrics and clade models in detecting those errors. The simulation was performed by swapping gene sequences between two species, one from inside and another from the species outside the clade of interest. The gene sequences were swapped in two ways. In non-reciprocal exchange, we replaced the selected gene’s sequences inside the clade with one from the outside clade. The species were selected randomly from both sides for this replacement. In the reciprocal exchange, gene sequences were swapped between two species, one from inside and another from outside the clade. One hundred datasets were generated for reciprocal and non-reciprocal swapping, which were then analyzed using *DrPhylo*.

## SUPPLEMENTARY FIGURES

**Supplementary Figure 1.**
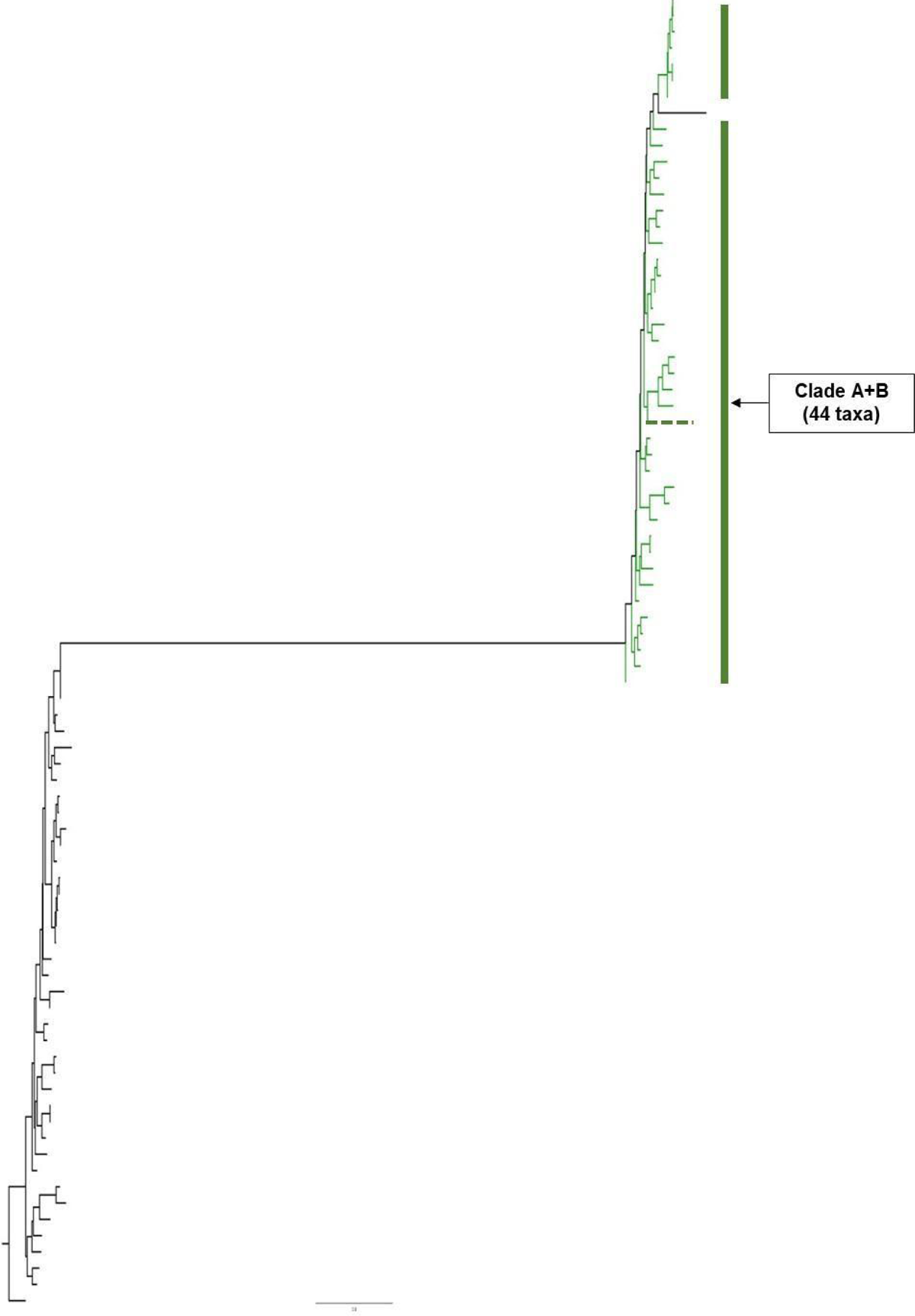
Gene tree of BUSCOfEOG7W9S51 (*7W9S51*). Species indicated by green tip branches belong to clade A+B in the phylogenomic tree (Fig. 2). The species forming clade A (*A. rubescence*) is marked by dotted green branch. The scale bar is in the units of the number of substitutions per site. The gene tree was inferred using the ML method and obtained from Shen et al. (2017).

**Supplementary Figure 2.**
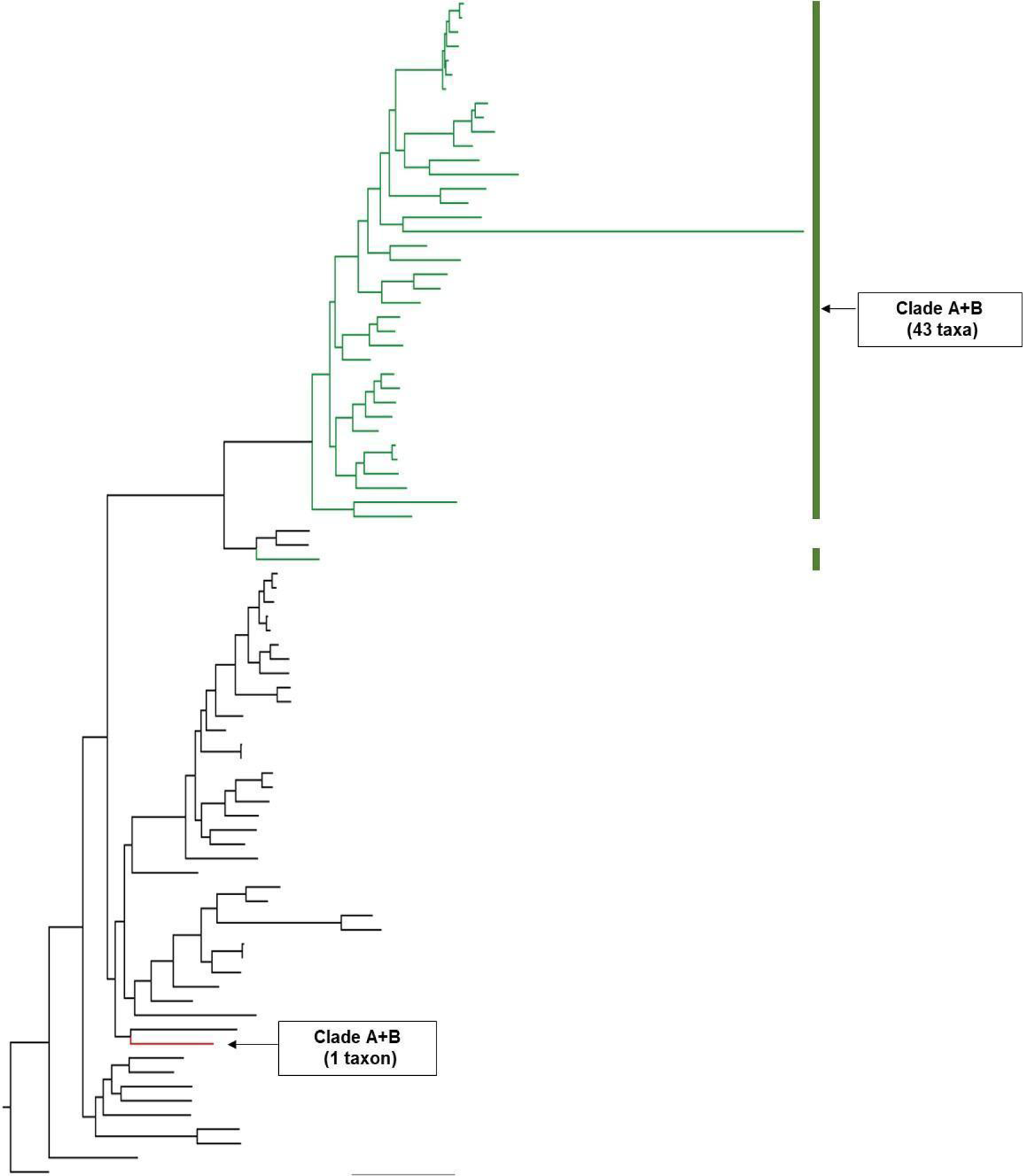
Gene tree of BUSCOfEOG7TN012 (*7TN012*). Species indicated by red and green tip branches lead to species belonging to clade A+B in the phylogenomic tree (see Fig. 2). Green tip branches led to species that are clustered with other species from the clade A+B (Fig. 2) and received a positive GSC score (Fig. 3b). On the other hand, the red tip branch is for the species (*A. rubescence*) from clade A+B, and received a negative GSC score for this gene. The scale bar is in the units of the number of substitutions per site. The gene tree was inferred by using the ML method and obtained from Shen et al. (2017).

**Supplementary Figure 3.**
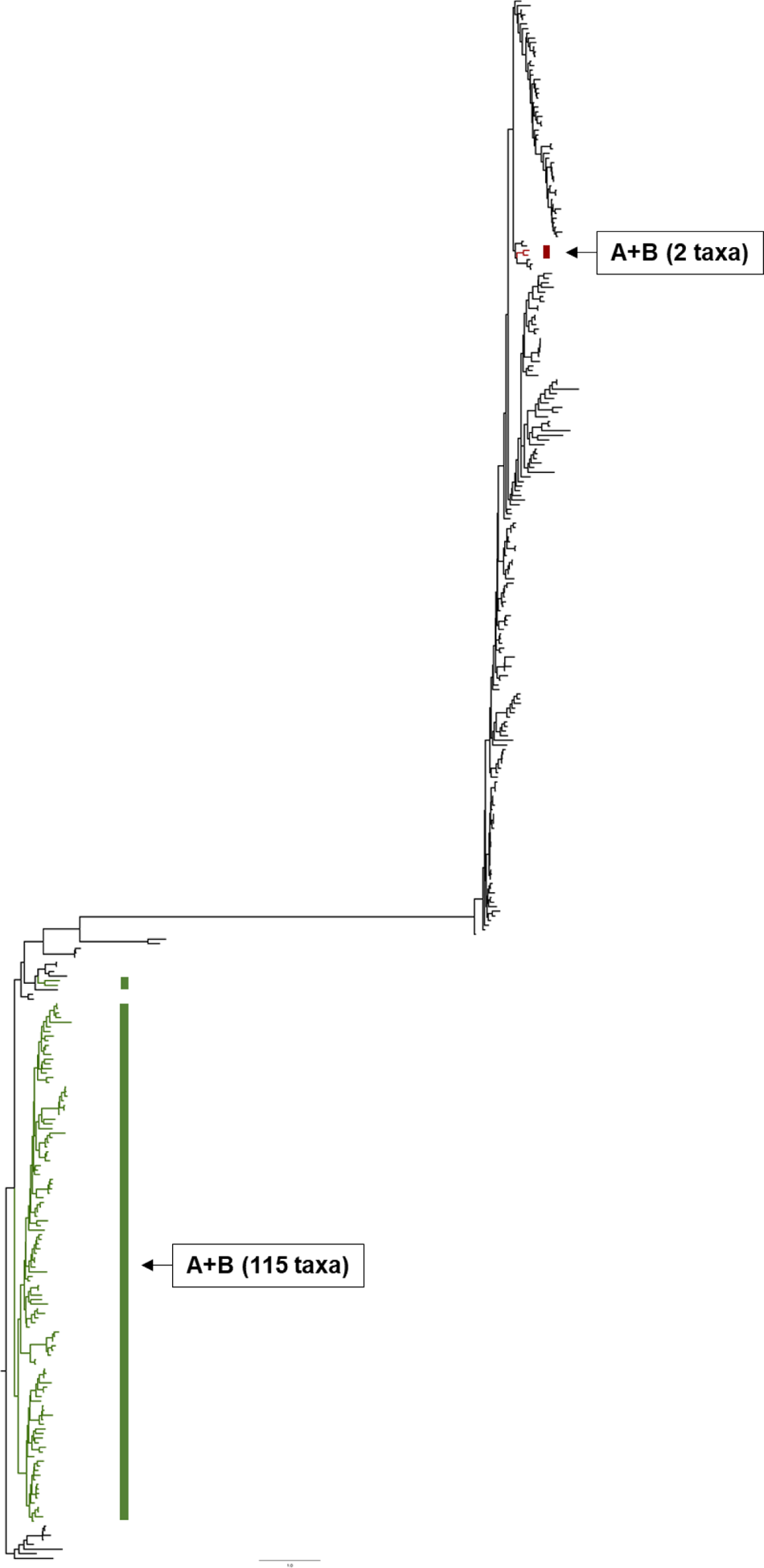
Gene tree of *EOG09343FGH*. Species indicated by red and green tip branches lead to species belonging to clade A+B in the phylogenomic tree (see Fig. 2). Green tip branches led to species that are clustered with other species from the clade A+B and received positive GSC scores (Fig. 4). On the other hand, species with tip branch negative GSC (Fig. 4) for this gene are separated from the clade A+B by an unexpectedly long internal branch. The scale bar is in the units of the number of substitutions per site. The gene tree was inferred by using the ML method and obtained from Shen et al. (2018).

**Supplementary Figure 4.**
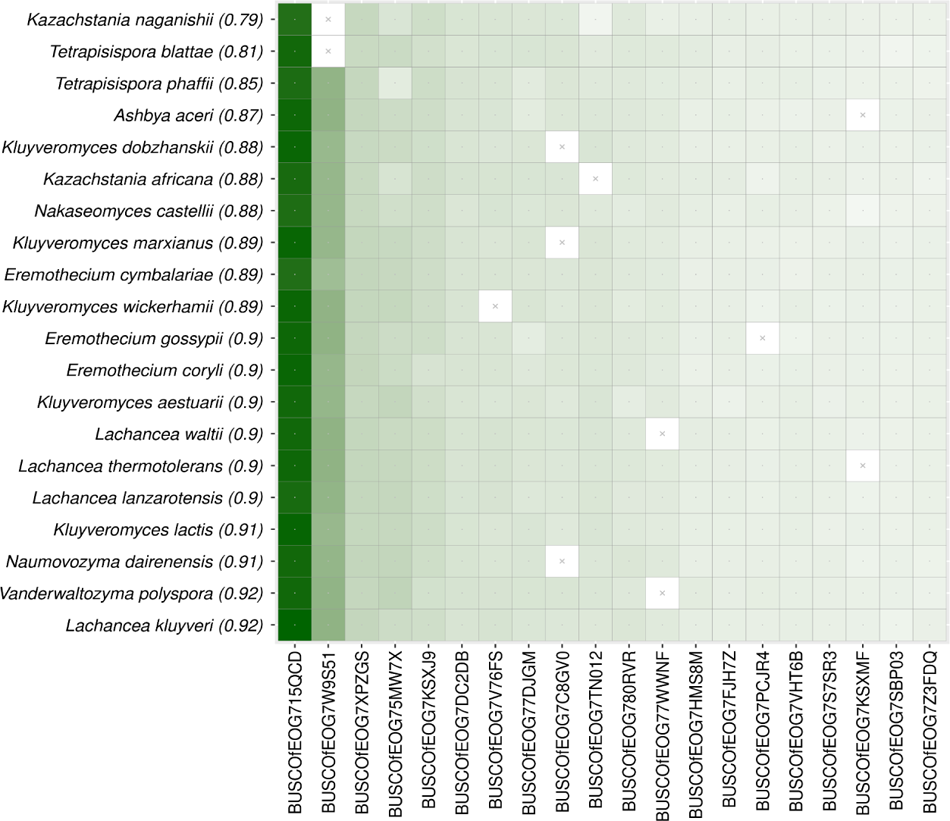
The M-grid for the control clade. The model grid for the *Saccharomycetaceae* clade. The M-grid shows that the gene *BUSCOfEOG715QCD* has the highest average *GSC* scores across all the taxa in the clade, whose CP is 0.79. *SCP* ranges from 1.00 to 0.79.

**Supplementary Figure 5.**
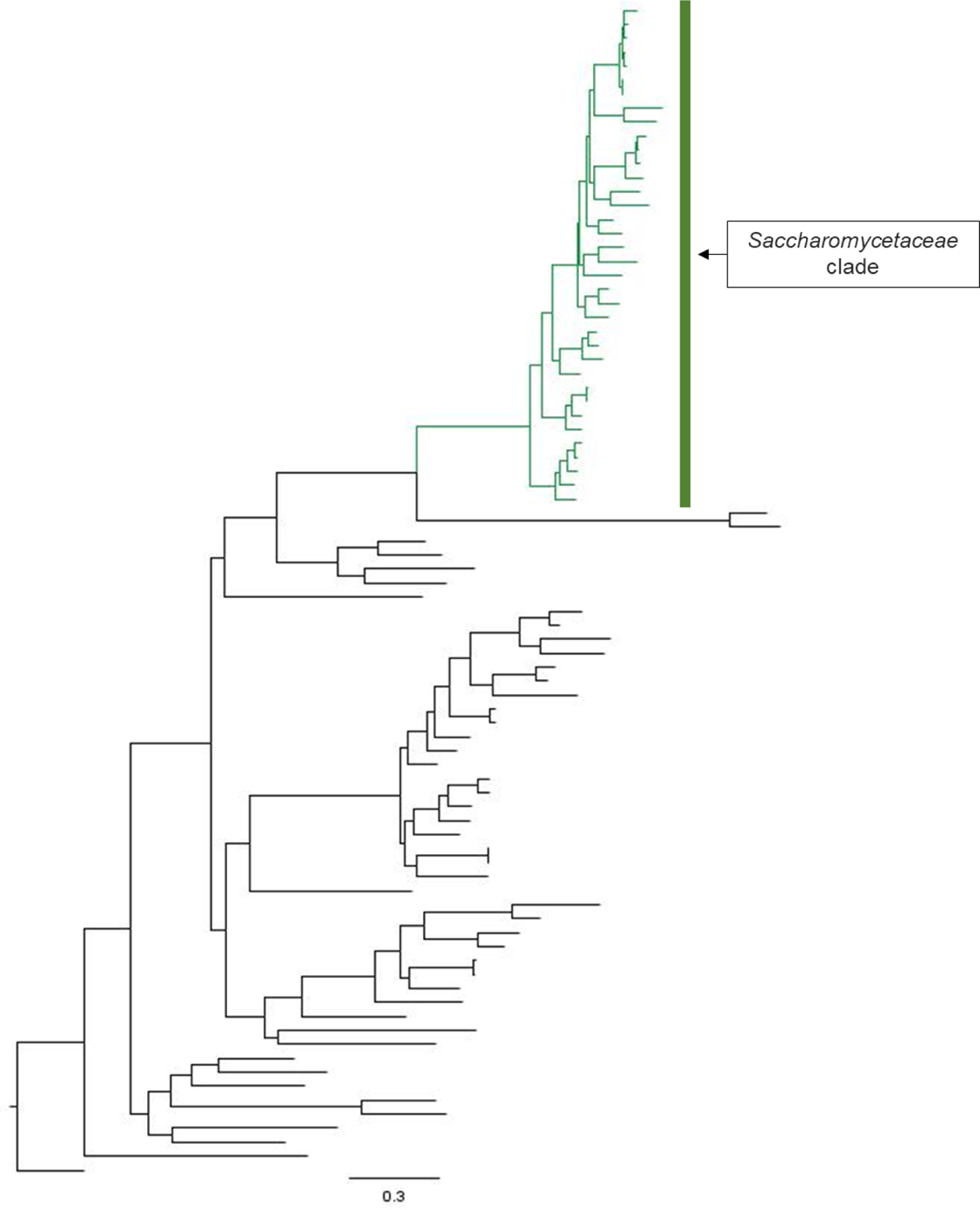
The ML phylogeny of *BUSCOfEOG715QCD* gene. Green branches mark tips belonging to species in *Saccharomycetaceae* clade and received positive *GSC* values (Supplementary Fig. 4). No species in the clade received a negative *GSC* value. The scale bar is in the units of the number of substitutions per site. The gene tree was inferred by using the ML method and obtained from Shen et al. (2017).

**Supplementary Figure 6.**
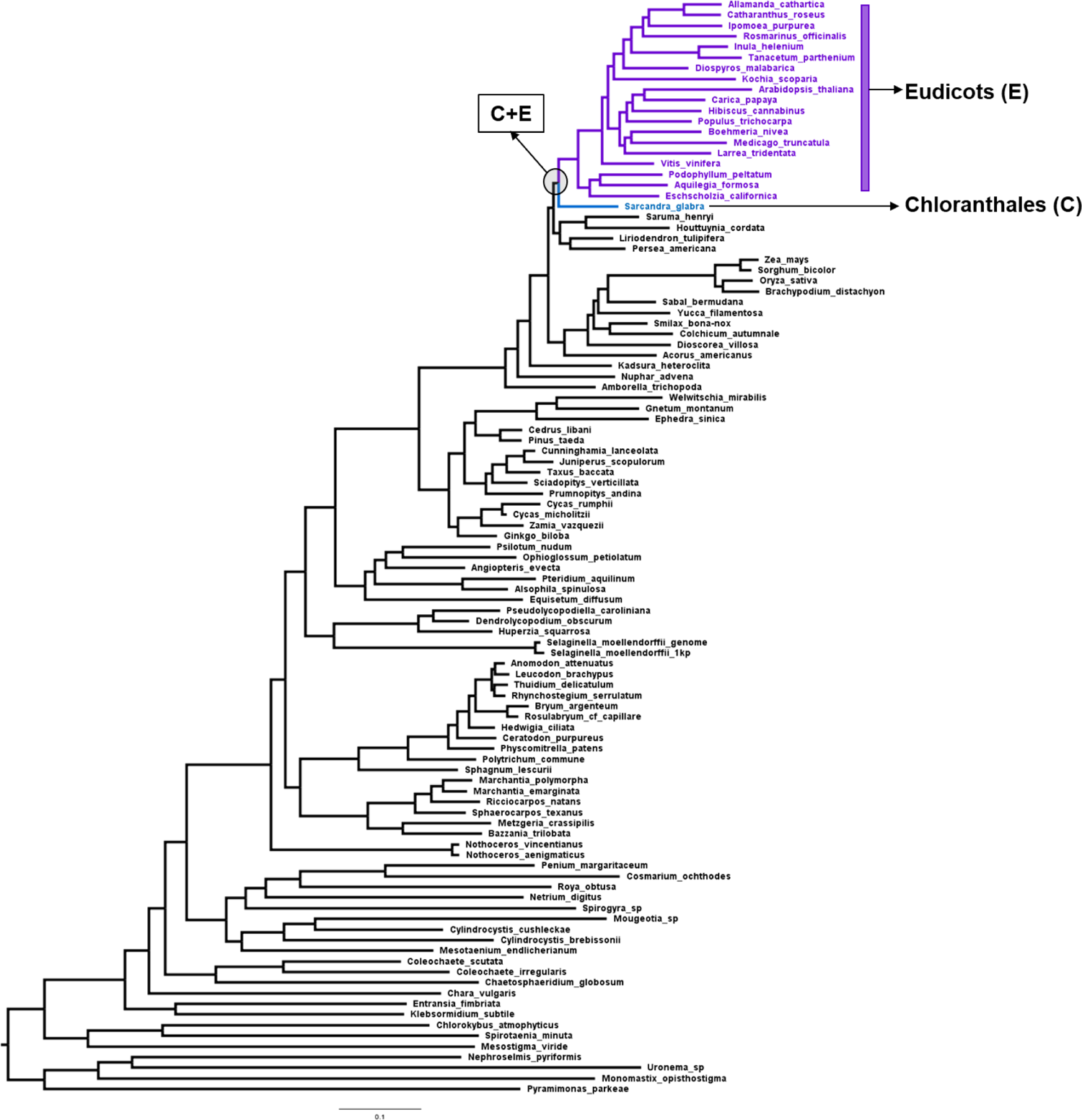
Plant phylogeny. The blue circle marks the clade containing 19 species of Eudicots (E; highlighted by purple-colored species) and one species from Chloranthales (C; highlighted by blue-colored species). The clade C+E was used to build the clade model. The highlighted blue branch leads to *Sarcandra glabra*, the only member of Chloranthales. The ML phylogenetic tree of 103 plant species inferred from the concatenated sequence alignment of 620 nuclear genes and obtained from Wickett et al. (2014).

**Supplementary Figure 7.**
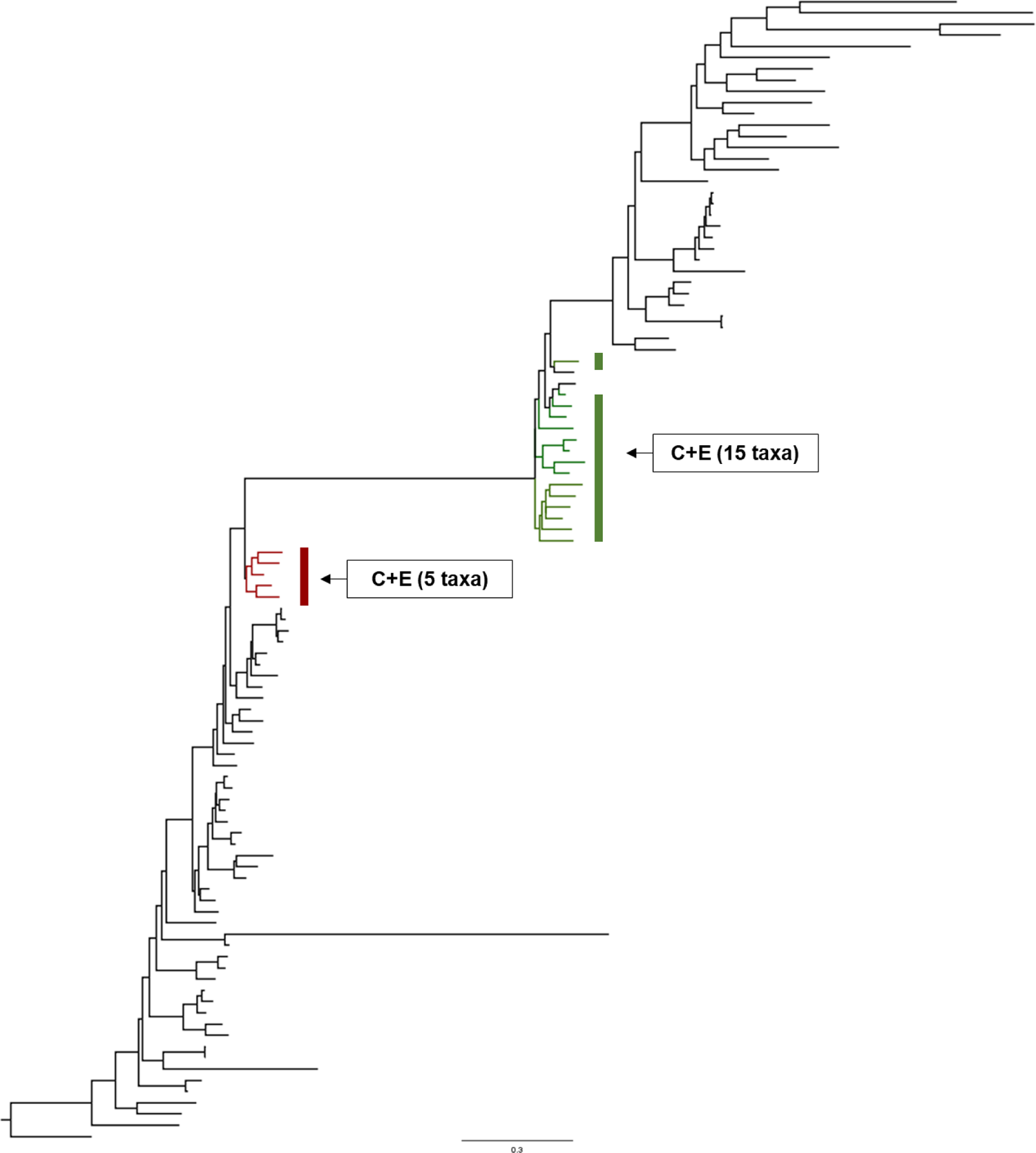
The ML phylogeny of 6040*_C12* gene. Green branches mark tips belonging to species in clade C+E and received positive GSC (Fig. 6). Red tips are for species that are separated from the majority of species in the C+E clade by an extremely long branch and receive negative GSC (Fig. 6) for this gene. The scale bar is in the units of the number of substitutions per site. The gene tree was inferred by using the ML method and obtained from Shen et al. (2017).

**Supplementary Figure 8.**
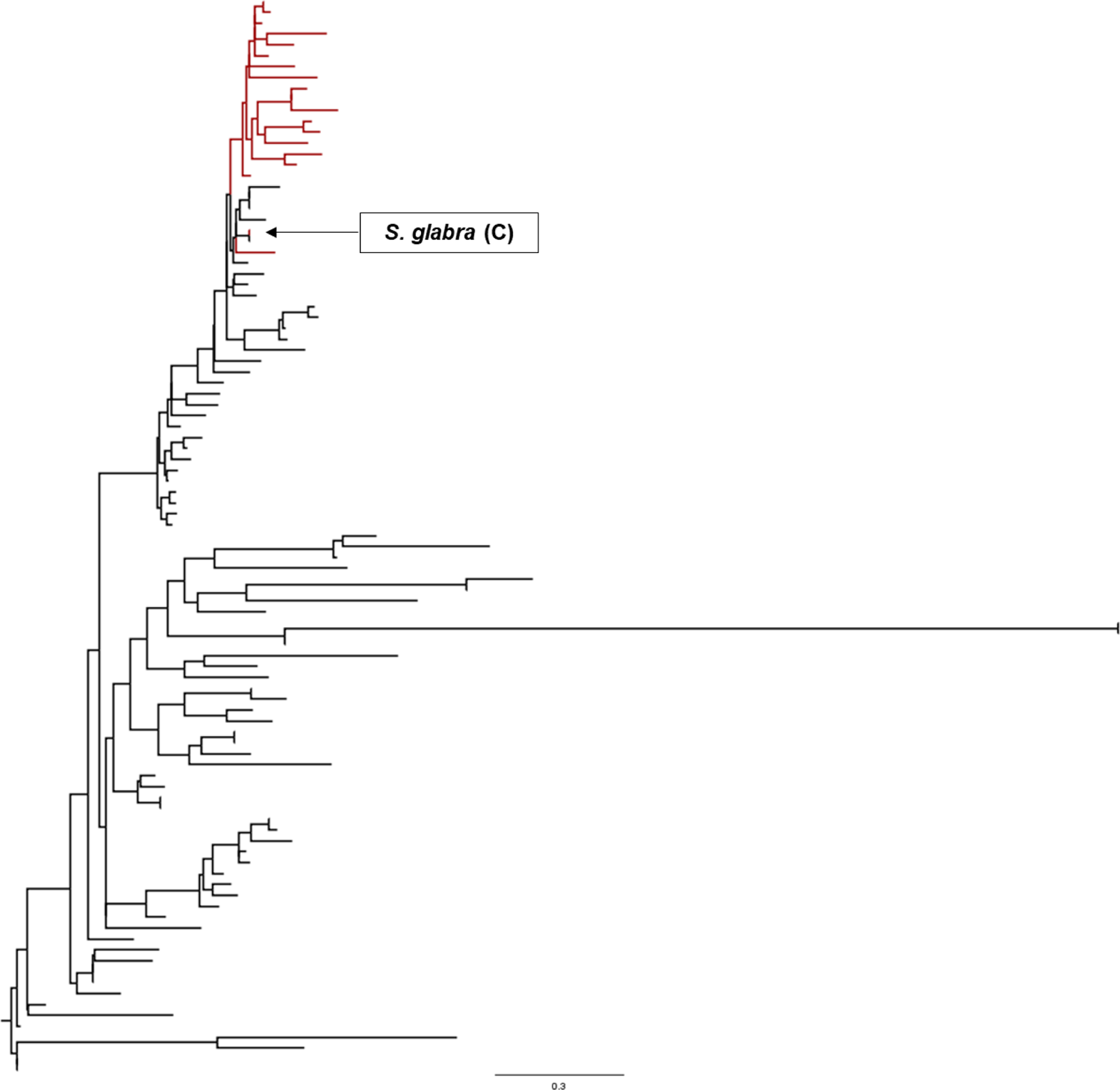
ML Gene tree of 5954*_C12*. Red branches mark tips belonging to species in clade C+E and receive negative GSC (Fig. 6). The clade (Chloranthales; C) has been indicated by an arrow containing one species, *Sarcandra glabra*. The scale bar is in the units of the number of substitutions per site. The gene tree was inferred by using the ML method and obtained from Shen et al. (2017).

**Supplementary Figure 9.**
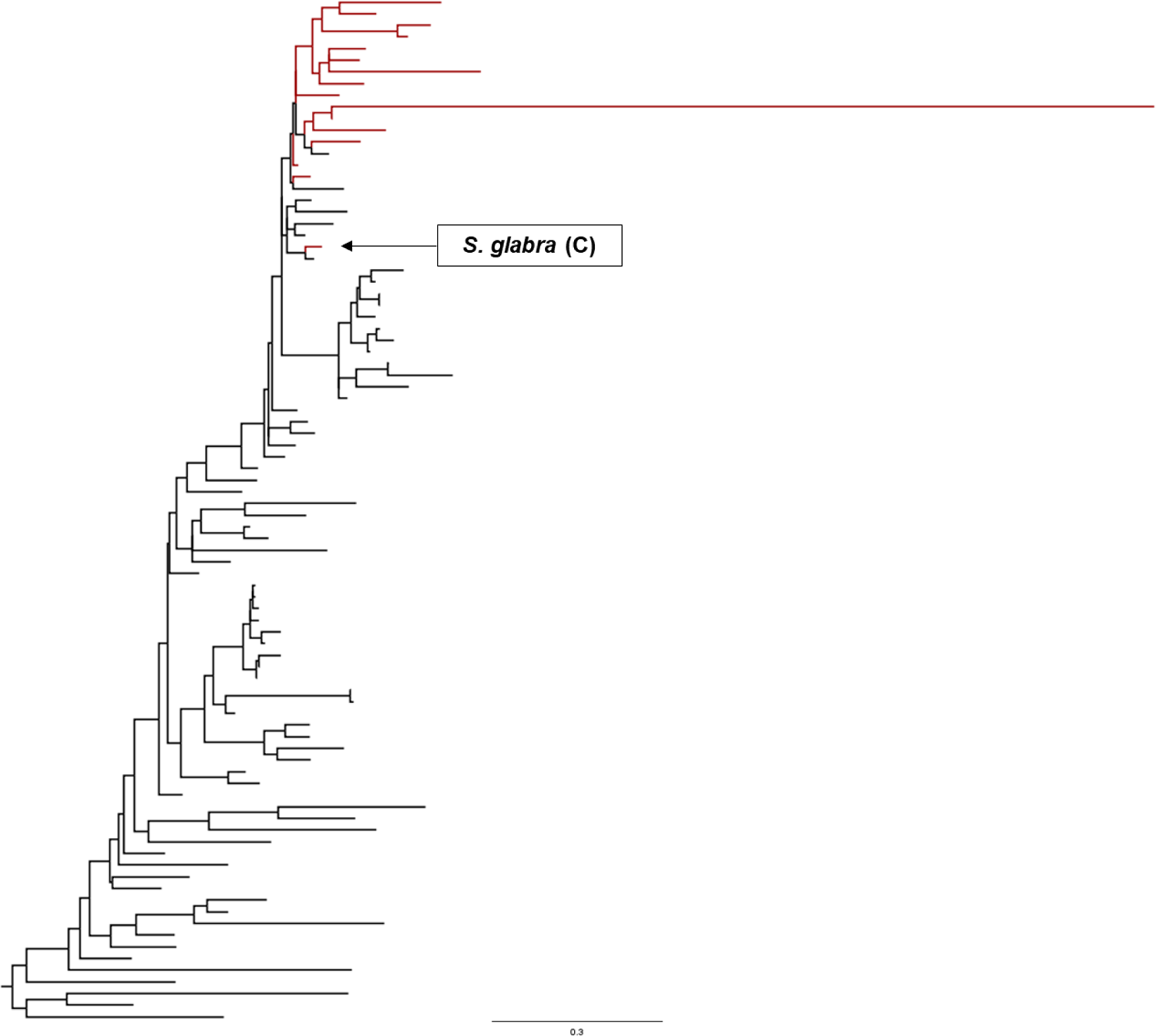
ML gene tree of 7684*_C12* Red branches mark tips belonging to species in clade C+E and receive negative GSC (Fig. 6). The clade (*Chloranthales*; C) has been indicated by an arrow containing one species, *Sarcandra glabra*. The scale bar is in the units of the number of substitutions per site. The gene tree was inferred by using the ML method and obtained from Shen et al. (2017).

**Supplementary Figure 10.**
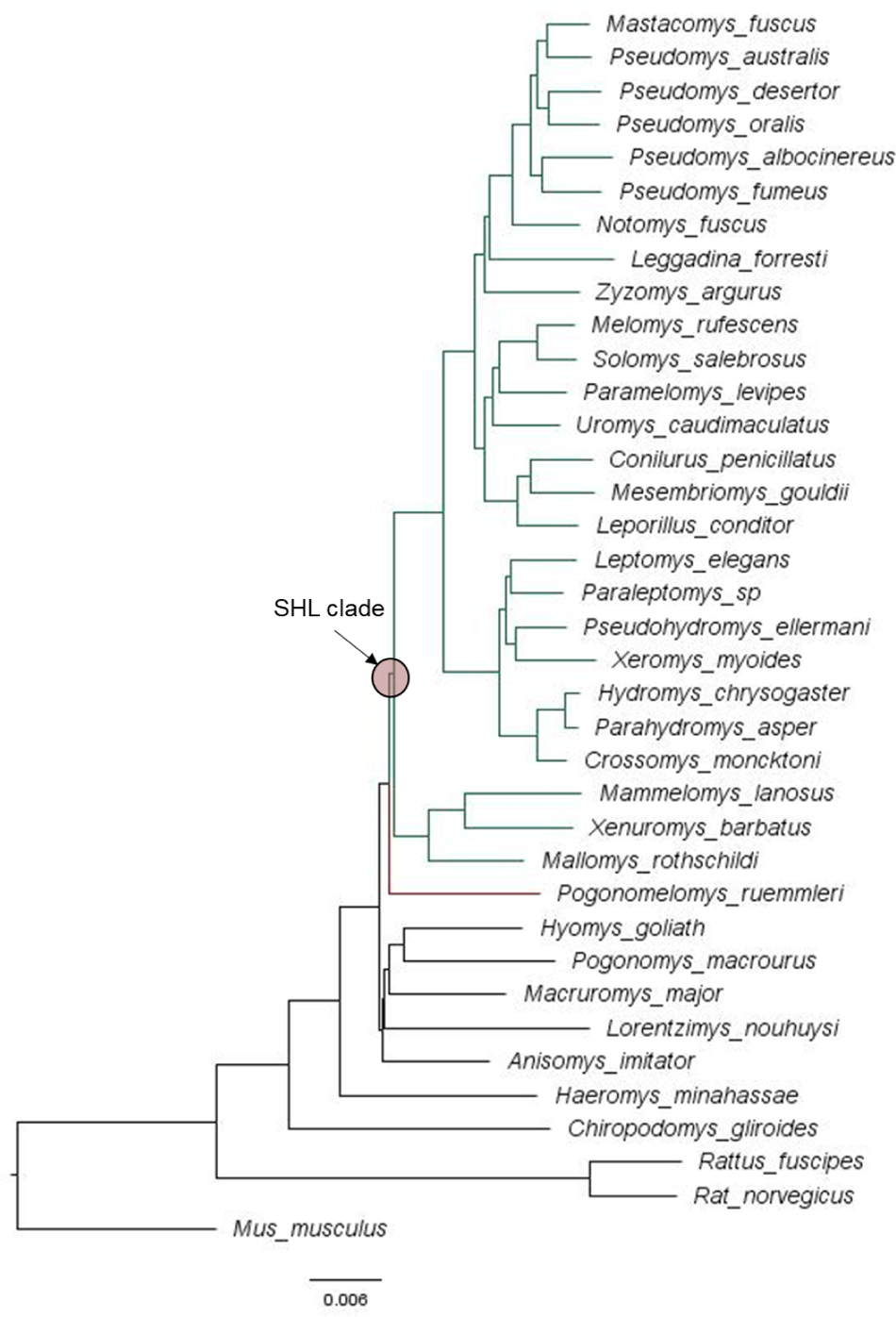
Animal phylogeny. The red circle marks the clade containing 26 species from the clade SHL. A total of 10 species were used to build the clade model. The highlighted red branch leads to *Pogonomelomys ruemmleri*, whose placement is different in trees inferred from the concatenated supermarix and MSC approaches. The ML phylogenetic tree of 37 rodents inferred from the concatenated sequence alignment of 1,245 nuclear genes and obtained from Roycroft et al. (2020).

**Supplementary Figure 11.**
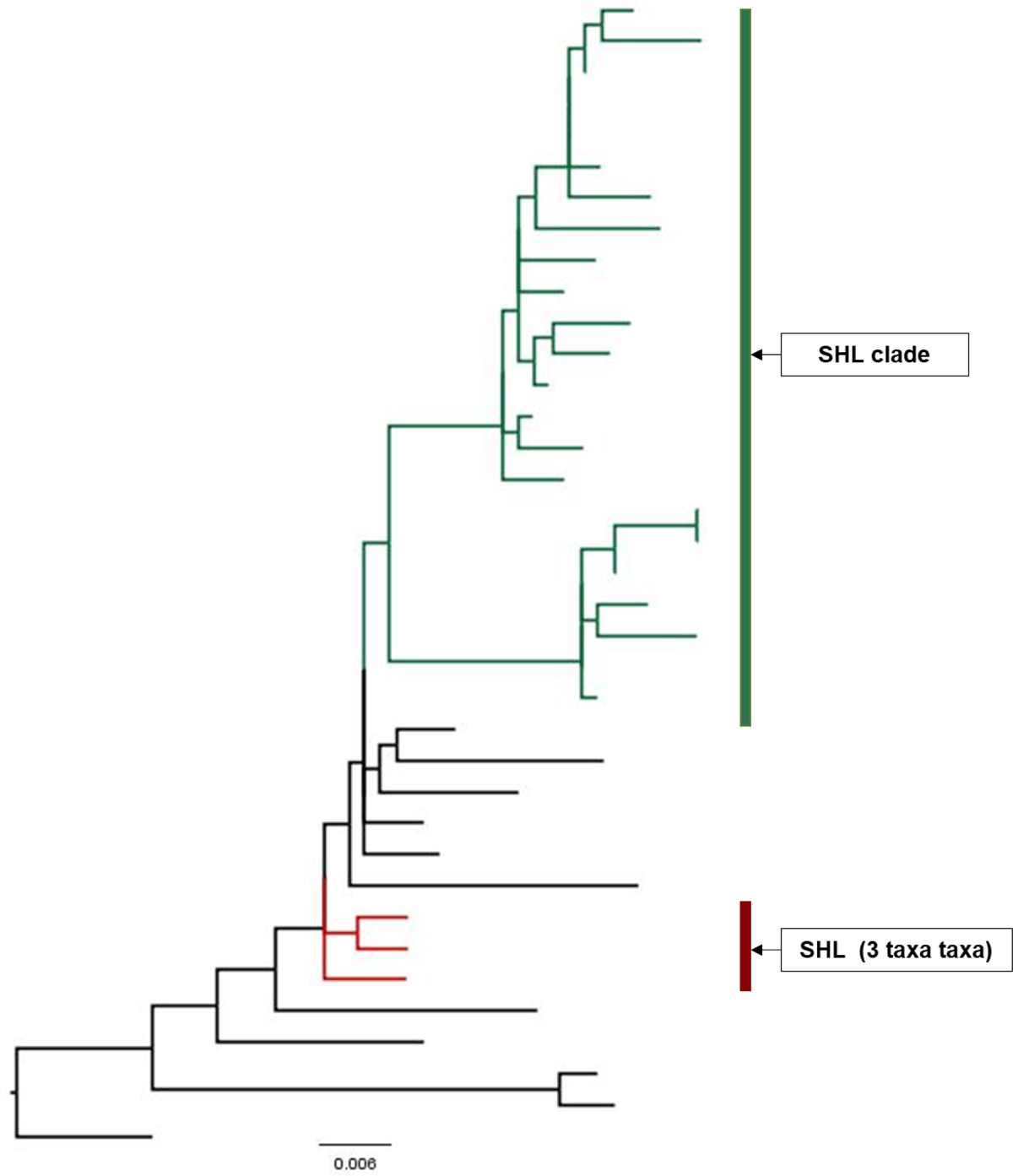
The ML phylogeny of *FOXJ1_1_rat*. Green branches mark tips belonging to species in clade SHL clade and received positive GSC (Fig. 7). Red tips are for species that are separated from the majority of species in the SHL clade and receive negative GSC (Fig. 7) for this gene. The scale bar is in the units of the number of substitutions per site. The gene tree was inferred using ML approach and obtained from Shen et al. (2021).

**Supplementary Figure 12.**
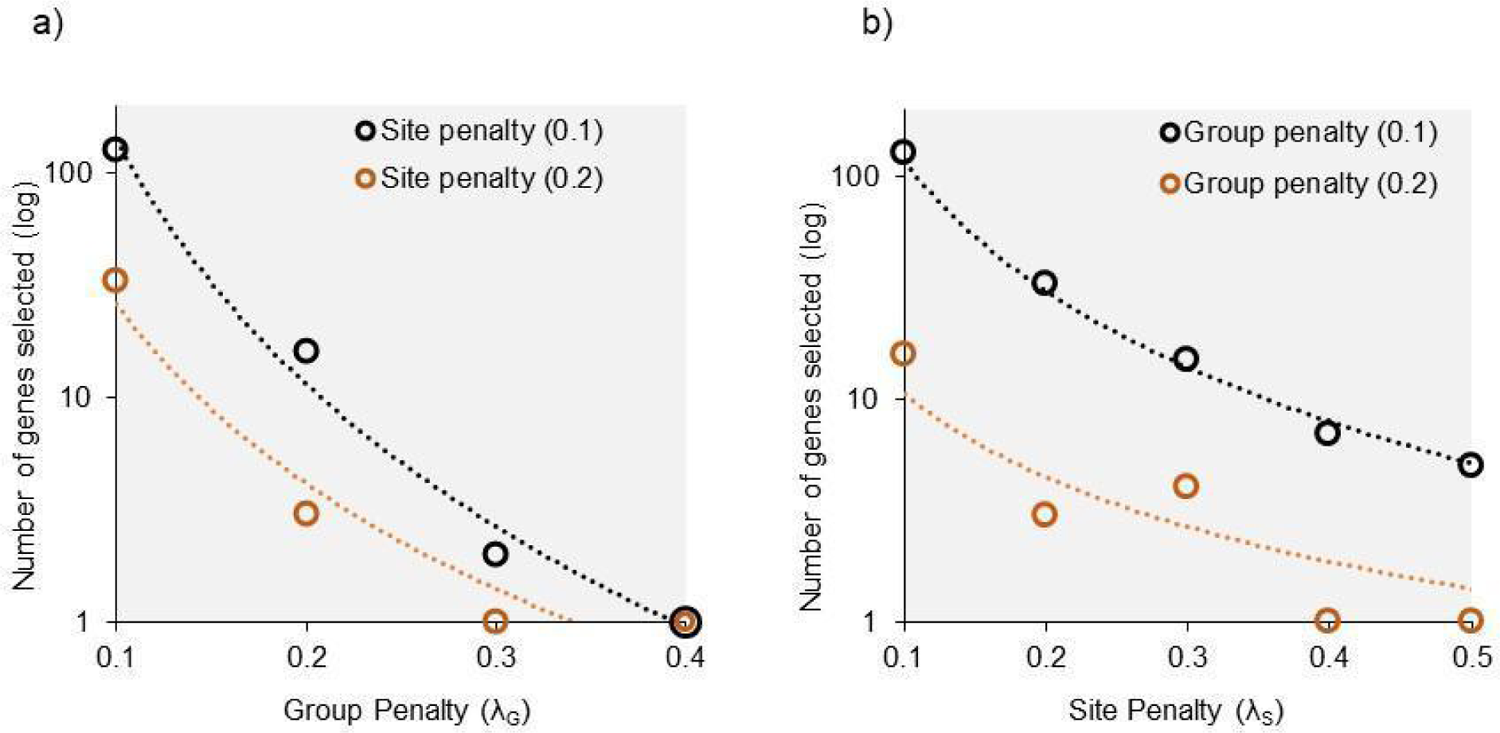
Relationship between the penalty parameters and the number of genes selected in the ESL model. (a) A scatterplot showing the relationship between the group penalty parameter (λ_G_) and the number of selected genes (log scale) in the model while keeping the site penalty fixed (λ_S_ = 0.1 and 0.2). As the group penalty parameter increases, the number of selected genes decreases exponentially, and only one gene is included in the clade model beyond a certain point (λ_G_ > 0.3). (b) A scatterplot depicting the influence of the site penalty parameter (λ_S_) on the number of selected genes in the model while holding the group penalty (λ_G_) fixed. The dotted lines show power curve fits (*R*^2^ > 0.8).

---

**Supplementary Table 1.**
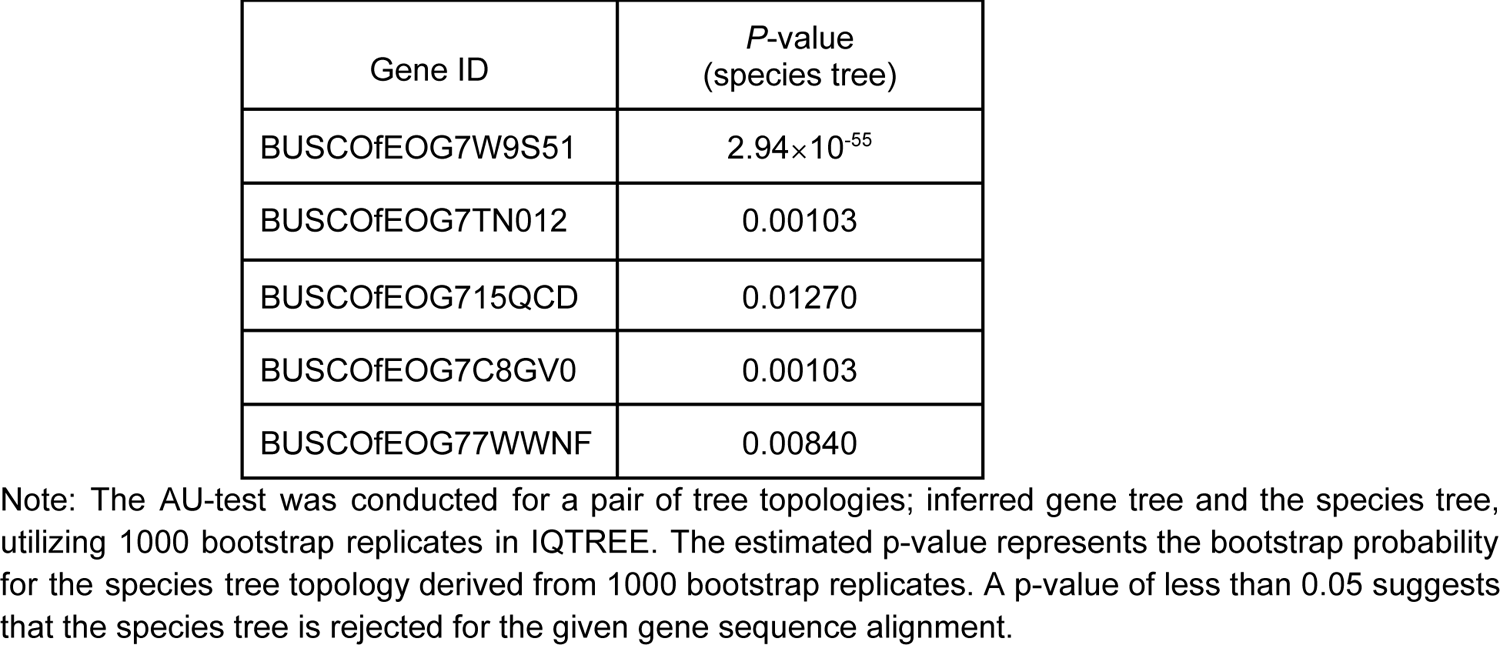
AU-test for candidate genes selected for A+B clade in fungi phylogeny.

| Gene ID | *P*-value<br>(species tree) |
| --- | --- |
| BUSCOfEOG7W9S51 | $2.94 \times 10^{-55}$ |
| BUSCOfEOG7TN012 | 0.00103 |
| BUSCOfEOG715QCD | 0.01270 |
| BUSCOfEOG7C8GV0 | 0.00103 |
| BUSCOfEOG77WWNF | 0.00840 |

Note: The AU-test was conducted for a pair of tree topologies; inferred gene tree and the species tree, utilizing 1000 bootstrap replicates in IQTREE. The estimated p-value represents the bootstrap probability for the species tree topology derived from 1000 bootstrap replicates. A p-value of less than 0.05 suggests that the species tree is rejected for the given gene sequence alignment.

**Supplementary Table 2.** Time taken (minutes) in *DrPhylo* analysis.

| Dataset (Type) | Clade | Number of taxa in the clade | Total number of genes | Genes selected in the model | Count of Models built (count of multi-gene models) | Total time (minutes) |
| --- | --- | --- | --- | --- | --- | --- |
| Fungus (AA) | A+B | 44 | 1,233 | 124 | 14 (12) | 29.5 |
| Fungus (AA) | Control | 36 | 1,233 | 244 | 48 (44) | 64.1 |
| Extended Fungus (AA) | A+B | 117 | 1,292 | 87 | 19 (14) | 51.8 |
| Plant (DNA) | C+E | 20 | 620 | 286 | 34 (33) | 7.9 |
| Animal (DNA) | SHL | 10 | 1,245 | 260 | 47(45) | 8.6 |

Note: Dataset names correspond to the species under analysis, while data type refers to the base type within each dataset (AA: Amino Acid). The clade ID is the identification code for the clades examined in the main text. We have also included the computational time required for comprehensive *DrPhylo* analysis for each clade. All the analyses were performed on a desktop computer with four cores and 16 GB of RAM.

## References

Abadi S, Azouri D, Pupko T, Mayrose I. 2019. Model selection may not be a mandatory step for phylogeny reconstruction. Nat. Commun. 10(1):1–11.

Brown JM, Thomson RC. 2016. Bayes factors unmask highly variable information content, bias, and extreme influence in phylogenomic analyses. Syst. Biol. 66(4):517–530.

Chiari Y, Cahais V, Galtier N, Delsuc F. 2012. Phylogenomic analyses support the position of turtles as the sister group of birds and crocodiles (Archosauria). Bmc Biol. 10(1):65.

Comte A, Tricou T, Tannier E, Joseph J, Siberchicot A, Penel S, Allio R, Delsuc F, Dray S, de Vienne DM. 2023. PhylteR: Efficient Identification of Outlier Sequences in Phylogenomic Datasets. Mol. Biol. Evol. 40(11).

Edwards S V. 2016. Phylogenomic subsampling: a brief review. Zool. Scr. 45:63–74.

Feuda R, Dohrmann M, Pett W, Philippe H, Rota-Stabelli O, Lartillot N, Wörheide G, Pisani D. 2017. Improved Modeling of Compositional Heterogeneity Supports Sponges as Sister to All Other Animals. Curr. Biol. 27(24):3864–3870.e4.

Fitzpatrick DA. 2012. Horizontal gene transfer in fungi. FEMS Microbiol. Lett. 329(1):1–8.

Freund Y, Schapire RE. 1999. Large margin classification using the perceptron algorithm. Mach. Learn. 37(3):277–296.

Gadagkar SR, Rosenberg MS, Kumar S. 2005. Inferring species phylogenies from multiple genes: Concatenated sequence tree versus consensus gene tree. J. Exp. Zool. Part B Mol. Dev. Evol. 304B(1):64–74.

Guimarães Fabreti L, Höhna S. 2023. Nucleotide Substitution Model Selection Is Not Necessary for Bayesian Inference of Phylogeny With Well-Behaved Priors. Syst. Biol.:syad041.

Hastie T, Tibshirani R, Wainwright M. 2015. Statistical learning with sparsity: The lasso and generalizations. CRC Press: Boca Raton, FL.

Hillis DM, Bull JJ. 1993. An empirical test of bootstrapping as a method for assessing confidence in phylogenetic analysis. Syst. Biol. 42(2):182–192.

Höhna S, Landis MJ, Huelsenbeck JP. 2021. Parallel power posterior analyses for fast computation of marginal likelihoods in phylogenetics. PeerJ 9:e12438.

Homziak NT, Storer CG, Gall LF, Borth RJ, Kawahara AY. 2023. Phylogenomics resolves major relationships of Catocala underwing moths. Syst. Entomol. 48(4):633–643.

Hughes LC, Nash CM, White WT, Westneat MW. 2023. Concordance and Discordance in the Phylogenomics of the Wrasses and Parrotfishes (Teleostei: Labridae). Syst. Biol. 72(3):530–543.

Jeffroy O, Brinkmann H, Delsuc F, Philippe H. 2006. Phylogenomics: the beginning of incongruence? Trends Genet. 22(4):225–231.

Kainer D, Lanfear R. 2015. The effects of partitioning on phylogenetic inference. Mol. Biol. Evol. 32(6):1611–1627.

Kapli P, Yang Z, Telford MJ. 2020. Phylogenetic tree building in the genomic age.Pellens R, Grandcolas P, editors. Nat. Rev. Genet. 21(7):428–444.

Kumar S. 2022. Embracing Green Computing in Molecular Phylogenetics. Mol. Biol. Evol. 39(3):43.

Kumar S, Filipski AJ, Battistuzzi FU, Kosakovsky Pond SL, Tamura K. 2012. Statistics and truth in phylogenomics. Mol. Biol. Evol. 29(2):457–472.

Kumar S, Sharma S. 2021. Evolutionary Sparse Learning for Phylogenomics. Mol. Biol. Evol. 38(11):4674–4682.

Lanfear R, Frandsen PB, Wright AM, Senfeld T, Calcott B. 2017. PartitionFinder 2: New Methods for Selecting Partitioned Models of Evolution for Molecular and Morphological Phylogenetic Analyses. Mol. Biol. Evol. 34(3):772–773.

Liu J, Ji S, Ye J. 2011. SLEP: Sparse learning with efficient projections. Note 6:491.

Liu J, Ye J. 2010. Moreau-Yosida regularization for grouped tree structure learning. In: Proceedings of the 23rd International Conference on Neural Information Processing Systems - Volume 2. Curran Associates Inc., NY. p. 1459–1467.

Liu K, Linder CR, Warnow T. 2011. RAxML and FastTree: Comparing two methods for large-scale maximum likelihood phylogeny estimation. PLoS One 6(11):e27731.

Lundberg SM, Erion G, Chen H, DeGrave A, Prutkin JM, Nair B, Katz R, Himmelfarb J, Bansal N, Lee SI. 2020. From local explanations to global understanding with explainable AI for trees. Nat. Mach. Intell. 2020 21 2(1):56–67.

Meier L, Van De Geer S, Bühlmann P. 2008. The group lasso for logistic regression. J. R. Stat. Soc. Ser. B Stat. Methodol. 70(1):53–71.

Mirarab S, Reaz R, Bayzid MS, Zimmermann T, S. Swenson M, Warnow T. 2014. {ASTRAL}: genome-scale coalescent-based species tree estimation. Bioinformatics 30(17):i541--i548.

Mongiardino Koch N. 2021. Phylogenomic Subsampling and the Search for Phylogenetically Reliable Loci. Mol. Biol. Evol. 38(9):4025–4038.

Nakhleh L. 2013. Computational approaches to species phylogeny inference and gene tree reconciliation. Trends Ecol. Evol. 28(12):719–728.

Philippe H, Delsuc F, Brinkmann H, Lartillot N. 2005. Phylogenomics. Annu. Rev. Ecol. Evol. Syst. 36(1):541–562.

Phillips MJ, Delsuc F, Penny D. 2004. Genome-Scale Phylogeny and the Detection of Systematic Biases. Mol. Biol. Evol. 21(7):1455–1458.

Phillips MJ, Delsuc FF, Penny D. 2004. Genome-Scale Phylogeny and the Detection of Systematic Biases. Mol. Biol. Evol. 21(7):1455–1458.

Redmond AK, McLysaght A. 2021. Evidence for sponges as sister to all other animals from partitioned phylogenomics with mixture models and recoding. Nat. Commun. 12(1).

Richards TA, Soanes DM, Foster PG, Leonard G, Thornton CR, Talbot NJ. 2009. Phylogenomic Analysis Demonstrates a Pattern of Rare and Ancient Horizontal Gene Transfer between Plants and Fungi. Plant Cell 21(7):1897–1911.

Riley R, Haridas S, Wolfe KH, Lopes MR, Hittinger CT, Göker M, Salamov AA, Wisecaver JH, Long TM, Calvey CH, et al. 2016. Comparative genomics of biotechnologically important yeasts. Proc. Natl. Acad. Sci. U. S. A. 113(35):9882–9887.

Rokas A, Williams BI, King N, Carroll SB. 2003. Genome-scale approaches to resolving incongruence in molecular phylogenies. Nature 425(6960):798–804.

Roycroft EJ, Moussalli A, Rowe KC. 2020. Phylogenomics Uncovers Confidence and Conflict in the Rapid Radiation of Australo-Papuan Rodents. Syst. Biol. 69(3):431–444.

Salichos L. 2014. Quantifying Phylogenetic Incongruence and Identifying Contributing Factors in a Yeast Model Clade.

Sanderford M, Sharma S, Tamura K, Stecher G, Liu J, Ji S, Ye J, Kumar S. 2024. myESL: A software for evolutionary sparse learning in molecular phylogenetics and evolutionary genomics. In preparation.

Schmitt I, Lumbsch HT. 2009. Ancient Horizontal Gene Transfer from Bacteria Enhances Biosynthetic Capabilities of Fungi. PLoS One 4(2):e4437.

Schrider DR, Kern AD. 2018. Supervised Machine Learning for Population Genetics: A New Paradigm. Trends Genet. 34(4):301–312.

Seo T-K. 2008. Calculating Bootstrap Probabilities of Phylogeny Using Multilocus Sequence Data. Mol. Biol. Evol. 25(5):960–971.

Shao Y, Zhou L, Li F, Zhao L, Zhang BL, Shao F, Chen JW, Chen CY, Bi X, Zhuang XL, et al. 2023. Phylogenomic analyses provide insights into primate evolution. Science (80-.). 380(6648):913–924.

Shen XX, Hittinger CT, Rokas A. 2017. Contentious relationships in phylogenomic studies can be driven by a handful of genes. *Nat*. Ecol. Evol. 2017 *15* 1(5):1–10.

Shen XX, Opulente DA, Kominek J, Zhou X, Steenwyk JL, Buh K V., Haase MABB, Wisecaver JH, Wang M, Doering DT, et al. 2018. Tempo and Mode of Genome Evolution in the Budding Yeast Subphylum. Cell 175(6):1533--1545.e20.

Shen XX, Steenwyk JL, Rokas A. 2021. Dissecting Incongruence between Concatenation- and Quartet-Based Approaches in Phylogenomic Data. Syst. Biol. 70(5):997–1014.

Shen XX, Zhou X, Kominek J, Kurtzman CP, Hittinger CT, Rokas A. 2016. Reconstructing the backbone of the saccharomycotina yeast phylogeny using genome-scale data. G3 Genes, Genomes, Genet. 6(12):3927–3939.

Smith SA, Moore MJ, Brown JW, Yang Y. 2015. Analysis of phylogenomic datasets reveals conflict, concordance, and gene duplications with examples from animals and plants. BMC Evol. Biol. 15(1):150.

Smith SA, Walker-Hale N, Walker JF. 2020. Intragenic conflict in phylogenomic data sets.Crandall K, editor. Mol. Biol. Evol. 37(11):3380–3388.

Song S, Liu L, Edwards S V., Wu S. 2012. Resolving conflict in eutherian mammal phylogeny using phylogenomics and the multispecies coalescent model. Proc. Natl. Acad. Sci. U. S. A. 109(37):14942–14947.

Steenwyk JL, Li Y, Zhou X, Shen X-X, Rokas A. 2023. Incongruence in the phylogenomics era. Nat. Rev. Genet. 2023:1–17.

Struck TH. 2014. Trespex-detection of misleading signal in phylogenetic reconstructions based on tree information. Evol. Bioinforma. 10:51–67.

Suvorov A, Hochuli J, Schrider DR. 2020. Accurate Inference of Tree Topologies from Multiple Sequence Alignments Using Deep Learning. Syst. Biol. 69(2):221–233.

Tao Q, Tamura K, Battistuzzi FU, Kumar S. 2019. A machine learning method for detecting autocorrelation of evolutionary rates in large phylogenies. Mol. Biol. Evol. 36(4):811–824.

Tibshirani R. 1996. Regression Shriknage and Selectino via the Lasso. J. R. Stat. Soc. Ser. B 58(1):267–288.

De Vienne DM, Ollier S, Aguileta G. 2012. Phylo-MCOA: A Fast and Efficient Method to Detect Outlier Genes and Species in Phylogenomics Using Multiple Co-inertia Analysis. Mol. Biol. Evol. 29(6):1587–1598.

Walker JF, Brown JW, Smith SA. 2018. Analyzing contentious relationships and outlier genes in phylogenomics. Syst. Biol. 67(5):916–924.

Wickett NJ, Mirarab S, Nguyen N, Warnow T, Carpenter E, Matasci N, Ayyampalayam S, Barker MS, Burleigh JG, Gitzendanner MA, et al. 2014. Phylotranscriptomic analysis of the origin and early diversification of land plants. Proc. Natl. Acad. Sci. U. S. A. 111(45):E4859–E4868.

Williams TA, Cox CJ, Foster PG, Szöllősi GJ, Embley TM. 2019. Phylogenomics provides robust support for a two-domains tree of life. *Nat*. Ecol. Evol. 2019 *41* 4(1):138–147

Yang Z. 1996. Maximum-likelihood models for combined analyses of multiple sequence data. J. Mol. Evol. 42(5):587–596.

Young AD, Gillung JP. 2020. Phylogenomics — principles, opportunities and pitfalls of big-data phylogenetics. Syst. Entomol. 45(2):225–247.

